# Molecular Parallelism Underlies Convergent Highland Adaptation of Maize Landraces

**DOI:** 10.1101/2020.07.31.227629

**Authors:** Li Wang, Emily B. Josephs, Kristin M. Lee, Lucas M. Roberts, Rubén Rellán-Álvarez, Jeffrey Ross-Ibarra, Matthew B. Hufford

**Affiliations:** Shenzhen Branch, Guangdong Laboratory for Lingnan Modern Agriculture, Genome Analysis Laboratory of the Ministry of Agriculture, Agricultural Genomics Institute at Shenzhen, Chinese Academy of Agricultural Sciences, Shenzhen, China; Department of Ecology, Evolution, and Organismal Biology, Iowa State University, Ames, Iowa, USA; Department of Evolution and Ecology, University of California Davis, Davis, CA, USA; Department of Plant Biology, Michigan State University, East Lansing, MI, USA; Langebio, Libramiento Norte Carretera Leon Km 9.6, 36821 Irapuato, Gto., Mexico; Department of Molecular and Structural Biochemistry, North Carolina State University, Raleigh, NC, USA; Genome Center and Center for Population Biology and Genome Center, University of California Davis, Davis, CA, USA

## Abstract

Convergent phenotypic evolution provides some of the strongest evidence for adaptation. However, the extent to which recurrent phenotypic adaptation has arisen via parallelism at the molecular level remains unresolved, as does the evolutionary origin of alleles underlying such adaptation. Here, we investigate genetic mechanisms of convergent highland adaptation in maize landrace populations and evaluate the genetic sources of recurrently selected alleles. Population branch excess statistics reveal strong evidence of parallel adaptation at the level of individual SNPs, genes and pathways in four independent highland maize populations, even though most SNPs show unique patterns of local adaptation. The majority of selected SNPs originated via migration from a single population, most likely in the Mesoamerican highlands. Polygenic adaptation analyses of quantitative traits reveal that alleles affecting flowering time are significantly associated with elevation, indicating the flowering time pathway was targeted by highland adaptation. In addition, repeatedly selected genes were significantly enriched in the flowering time pathway, indicating their significance in adapting to highland conditions. Overall, our study system represents a promising model to study convergent evolution in plants with potential applications to crop adaptation across environmental gradients.

## Introduction

Convergent adaptation of populations to similar environments provides compelling evidence that natural selection, not neutral processes, shape trait variation. Similar phenotypes often arise independently in multiple species or populations exposed to the same evolutionary pressure [1, 2]. For example, species from four orders of insects, spanning 300 million years of divergence, have independently evolved tolerance to toxic compounds in milkweed and dogbane plant species [3, 4]. Likewise, reduction-of-function alleles at the FLC focus have evolved multiple times independently in *Capsella rubella* populations, conferring variation in flowering time [5]. Pool et al.[6] investigated cold adaptation across three pairs of highland and lowland *Drosophila melanogaster* populations, finding strong evidence for alleles that were repeatedly selected during highland colonization.

But while convergent phenotypes have been observed across the tree of life, in many cases their underlying genetic basis (*e.g*., molecular parallelism in which the same nucleotides, genes, or pathways are targeted versus convergence through independent molecular means) is unknown [7, 8]. Study of the genetics of convergence can help shed light on fundamental questions in evolutionary biology, including whether natural selection is constrained and repeatable or instead characterized by many molecular paths to similar phenotypes. The answer to these questions may depend, to a certain extent, on the trait itself. For example, the genetic architecture of a phenotype is a primary determinant of whether trait convergence results from parallel selection on orthologous loci [9]. Such convergence is more likely for simple traits where only a few loci contribute or when loci are subject to antagonistic pleiotropy and less likely for highly polygenic traits such as biomass or height [7, 9, 10].

Convergence through molecular parallelism, when it occurs, can originate in a number of ways. Genetic variation underlying convergent traits can arise as independent mutations, be derived from existing standing variation in a shared ancestral population, or be transferred between populations via migration [11]. Empirical studies have documented each of these modes of parallel adaptation. For instance, independent evolution of C4 photosynthesis in grasses [12] and sedges [13] has involved multiple *de novo* mutations in the phosphoenolpyruvate carboxylase (PEPC) gene, a key C4 enzyme. In contrast, parallel adaptation in rabbit populations resistant to the myxoma virus in Australia, France, and the United Kingdom is achieved through selection on standing variation in immunity-related genes [2]. Finally, convergence in patterns of wing coloration in butterfly species, an example of Müllerian mimicry, has resulted from gene flow across species [14, 15].

Given its history, maize is an ideal model system to study parallel adaptation. Maize was first domesticated in the warm lowlands of the Balsas River Valley approximately 9,000 years ago [16, 17], and subsequently spread to several independent highland regions, first to the the Mexican Central Plateau, and then to the highlands of the southwestern United States, Guatemala and the Andes [18, 19, 20, 21, 22, 23, 24, 25, 26, 27, 28, 29]. While highland regions colonized by maize are far from identical, commonalities include a shorter growing season, low temperature, low partial pressure of atmospheric gases, and high ultraviolet radiation [30, 31]. Highland individuals exhibit several traits that are thought to be adaptive under these conditions, including highly pigmented and hairy leaf sheaths [32]. Common garden experiments have also demonstrated that highland maize flowers substantially earlier than lowland material [33, 34, 35].

The genetic basis of highland phenotypes across these populations is largely unknown, however. Earlier population genetic analysis of maize in Mexico and the Andes identified a small percent of SNPs showing evidence of selection in both populations, but concluded that maize highland adaptation arose largely independently in the two populations [36]. One contributor to the differences between these populations has been gene flow from wild relatives: adaptive introgression from the wild congener teosinte (*Zea mays* ssp. *mexicana*, hereafter *mexicana*) is thought to have played an important role during highland adaptation in Mexico [37], but maize has no wild relatives in South America and simulations suggested that long-distance gene flow between the populations is unlikely [36]. Nonetheless, this earlier work was limited to two highland populations and relied on limited genotyping data. Subsequent investigation using whole genome sequencing, for example, has discovered alleles introgressed from teosinte in both the Guatemalan and southwestern US highlands outside of the distribution of *mexicana* [38], suggesting that migration from the Mexican highlands may not be as implausible as previously thought.

Here, we evaluate evidence for trait convergence through parallel molecular means using resequencing data from four highland populations of maize (Southwestern US, Mexico, Guatemala and the Andes) and two lowland populations (Mexico and South America). We assess the prevalence of molecular parallelism at the SNP, gene, and pathway levels, comparing models of *de novo* mutation, standing variation, and gene flow. Finally, our analysis of parallelism in pathways focuses on the well-characterized flowering time genes in maize and assesses signatures of polygenic adaptation for this trait.

## Results

### Description of samples and data

Four independent highland populations (the Southwestern US, Mexico, Guatemala and the Andes) and two reference lowland populations (Mexico and South America) were sampled to investigate highland adaptation in maize landraces (fig. 1). SNPs called from high-depth whole genome re-sequencing data were obtained from our previous study [38]. After filtering (see Materials and Methods), a total of 1,567,351 SNPs across 35 samples were retained for further analyses. We also characterized how environmental factors varied across our highland and lowland populations. A principal component analysis of 19 bioclim environmental variables (http://www.worldclim.org/) revealed that highland and lowland accessions were differentiated along PC1 (comprising 51.8% of variation); while PC2 (comprising 22.8% of variation) reflected latitudinal differences across highland populations (supplementary fig. S1, Supplementary Material online). By plotting the loadings from the 19 bioclim variables, we found PC1 was primarily polarized by temperature seasonality and PC2 by precipitation (supplementary fig. S1, Supplementary Material online).

**Figure 1:**
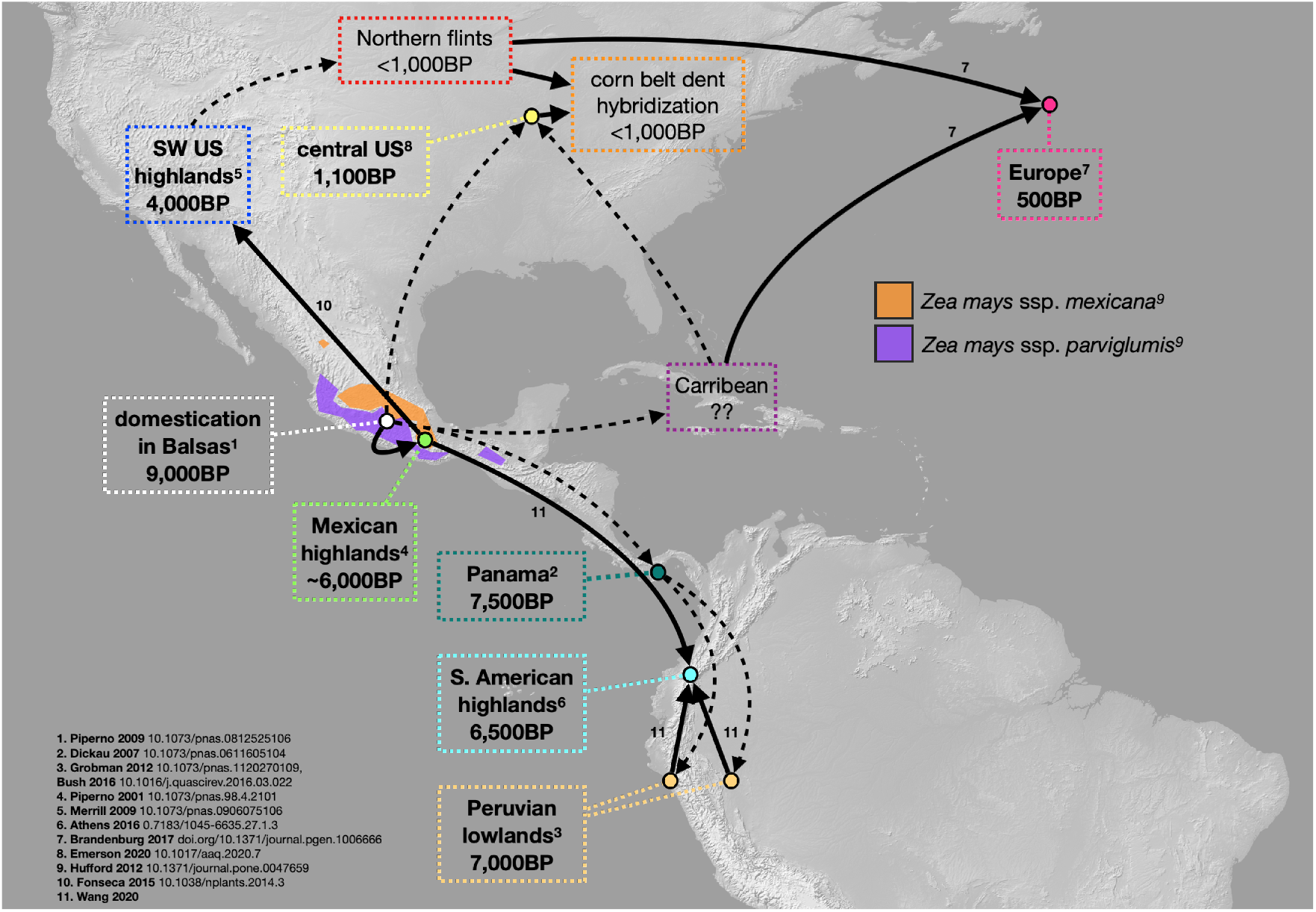
Sampling locations and expansion route of maize landraces. Domestication and expansion times for maize populations are from published articles [18, 19, 20, 21, 22, 23, 24, 25, 26, 27, 28, 29].

### Convergence through parallelism at the SNP level

In order to detect loci targeted by selection in highland populations, we utilized the Population Branch Excess (*PBE*) statistic [6], which characterizes changes in allele frequencies in a focal population relative to two independent “outgroup” populations [6]. *PBE* values were calculated for each SNP site and the top 5% were considered outliers and potential targets of selection.

To gauge the extent of convergence across our highland populations at the SNP level, we evaluated whether shared *PBE* outliers in pairs of populations and larger groups were more common than expected by chance. Significant overlap was observed in all pairs of highland populations (*p* ≈ 0, hypergeometric statistic test; supplementary table S1; supplementary table S2, Supplementary Material online). The Mexican Highland population showed the most substantial overlap in outlier SNPs with the Guatemalan Highlands (7.5-fold enrichment) and the Southwestern US Highlands (6-fold enrichment). While significantly more than expected, shared outliers between the Andes and all other highland populations were not nearly as extensive (between 2- and 4-fold enrichment), consistent with reduced parallelism at the SNP level between these isolated regions. The intersection of outlier SNPs across all highland populations was also significantly enriched (supplementary fig. S2, Supplementary Material online). Enrichment of overlapping selected sites was also confirmed in a more stringent tail of the top 1% of PBE values with qualitatively similar results (supplementary fig. S3, Supplementary Material online). While we did observe significant enrichment for parallelism at the SNP level, it is important to note that the majority of selected SNPs (from 59.7% to 68.1%) showed signatures of adaptation in a single highland population.

Our analysis of convergence at the SNP level using PBE revealed some particularly compelling candidate loci. For example, a SNP within PIF3.1 (phytochromeinteracting factor) exhibited the highest *PBE* value in the Mexican Highland population and was also detected as a target of selection in all other highland populations (fig. 2A). A non-synonymous, derived allele at this locus is fixed across all highland populations (fig. 2C). In addition, SNPs within GRMZM2G078118, a gene included in the jasmonic acid biosynthesis pathway, were detected as outliers in all Mesoamerican highland populations (SW US, Mexican Central Plateau, Guatemalan Highlands), but not the Andes (fig. 2BD).

**Figure 2:**
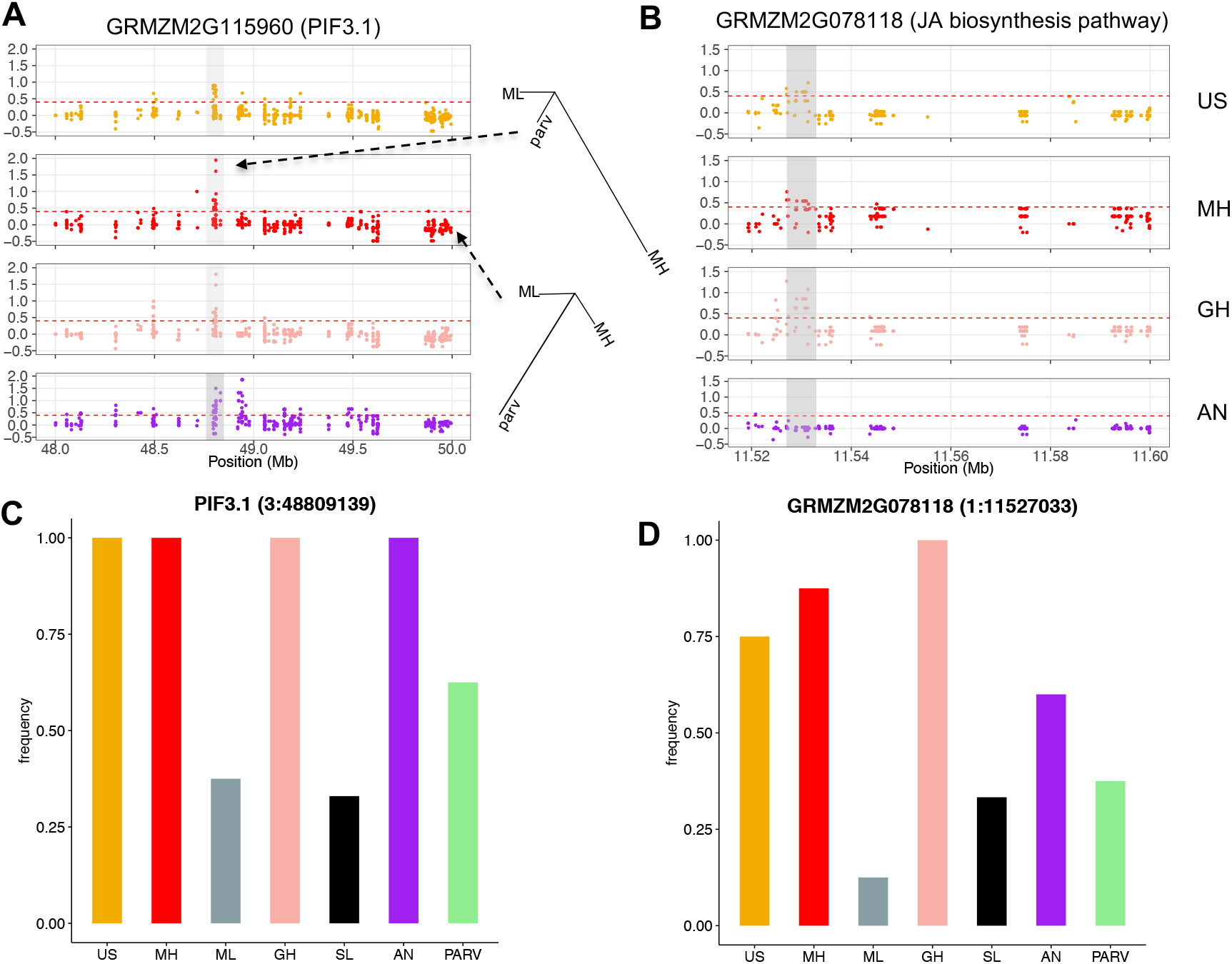
Patterns of parallel adaptation at two candidate loci. A. Distribution of *PBE* values in a 2-Mb region around the gene PIF3.1. The branch length of inset trees was based on *PBE* values and indicates the difference between a selected and unselected SNP. B. Distribution of *PBE* values in a 0.5 Mb region around the gene GRMZM2G078118 involved in jasmonic acid biosynthesis. C. Barplot of the reference allele frequency of one SNP located in PIF3.1. D. Barplot of the non-reference allele frequency of one SNP located in GRMZM2G078118.

While our PBE analysis reveals that selection targeted many of the same SNPs for adaptation across highland populations, this test alone does not clarify whether the same specific alleles were adaptive. We term outlier SNPs with the same allele elevated to high frequency in highland population pairs “co-directional”, and SNPs with different alleles as “anti-directional”. Predominantly, shared outliers in highland population pairs show co-directional allele frequency shifts (> 95% in all population pairs with the exception of the Southwestern US/Andes comparison, which was 87.6%). These results provide stronger evidence of molecular parallelism at the SNP level (fig. 3A). In contrast, random samples of SNPs that do not show evidence of selection exhibit much reduced signals of co-directional change. While random SNP samples between the Andes and other highland populations show roughly equal proportions of co- and anti-directionality, substantially more co-directional SNPs are observed in comparisons of Mesoamerican populations, perhaps reflecting their more closely shared histories (fig. 3A). However, the strong excess of co-directional changes in candidate SNPs may also be influenced by ancestral frequencies in lowland populations. For example, neutral drift has a greater likelihood of simultaneously fixing the same allele in multiple highland populations if the allele is already at high frequency in the lowlands, or, similarly, removing alleles at low frequency in the lowlands. In order to address this bias, we first approximated the ancestral allele frequencies of outlier SNPs (based on the twodimensional site frequency spectrum of Mexican lowland maize and the wild relative *Zea mays* ssp. *parviglumis*) in a subset of the neutral SNP set, and recalculated the number of co-directional and anti-directional SNPs. The same pattern was revealed (supplementary fig. S4, Supplementary Material online). Second, we subsampled outlier SNPs with a high minor allele frequency (*i.e*., between 0.3 and 0.5) and re-calculated the ratio of co-directional versus anti-directional SNPs. We found anti-directional SNPs were slightly increased in this subset, but co-directional SNPs were still far more numerous (fig. 3A). Taken together, the excess of co-directional SNPs in the outlier set was not biased by ancestral allele frequencies and the directionality of allele frequency changes in shared outlier SNPs provides strong evidence of molecular parallelism at the SNP level among highland maize populations.

**Figure 3:**
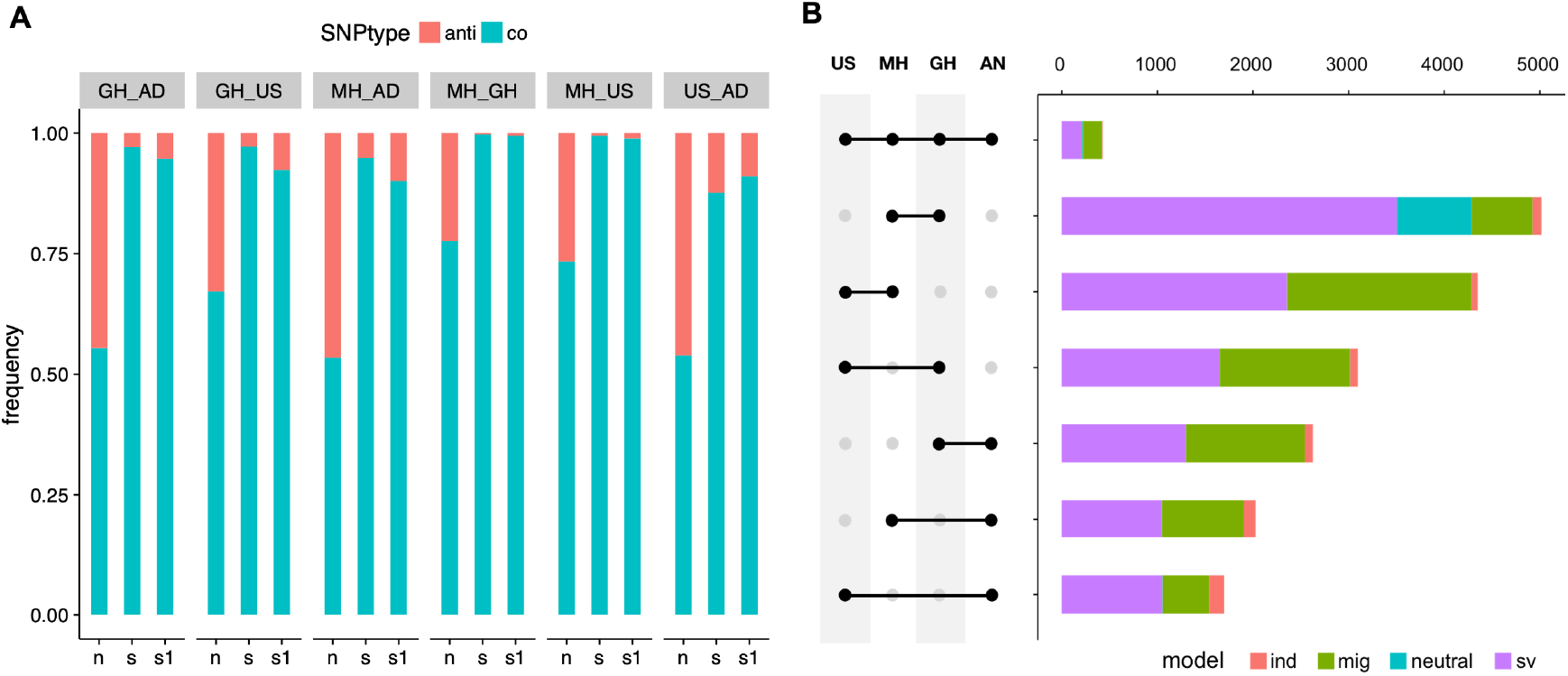
Patterns of parallel adaptation across loci. A. Distribution of co-directional and anti-directional SNPs in neutral and outlier SNP sets. n: neutral SNPs; s: common outliers; s1: a subset of common outliers with MAF between 0.3 – 0.5. B. Number of best fit models for genetic source of repeatedly selected SNPs. ind: independent *de-novo* mutation; mig: migration; sv: standing variation. Abbreviations for populations: AN, Andes; GH, Guatemalan Highlands; MH, Mexican Highlands; US, Southwestern US Highlands.

#### The source of shared adaptation among highland populations

To determine whether shared *PBE* outlier SNPs had independent origins in highland populations, we applied DMC, a composite-likelihood-based inference method that distinguishes among possible mechanisms [11]. We controlled for possible effects of linkage disequilibrium by subsampling outliers to one SNP per 2-kb window. First, we examined SNPs selected in all four independent highland populations. We found the vast majority of these common outliers (408 among 428) had the highest composite likelihood under models where there was a single origin of the beneficial allele. Within this subset, 49.1% and 46.3% of outliers had the standing variant model from a source population and the migration model as best fits, respectively (fig. 3B). The standing variant model from a source population (outlined in detail in Appendix A.4 of [11]) specifies that a beneficial allele originates in a single source population where it spreads via gene flow to other adapted populations where it may segregate for a standing time *t* prior to the onset of selection. For the common outliers identified under the migration model, the main migration sources were the Mexican (43.9%) and Guatemalan (41.9%) Highland populations. This result suggests that, for many shared outlier SNPs, gene flow among highland populations may have been an important source of adaptive variation.

We further explored the source of shared adaptive variation across pairwise samples of highland populations. As seen in four-wise comparisons, the standing variant model from a source population (49.4% to 69.9%; mean 56.7%) and the migration model (12.7% to 47.4%; mean 36.6%) were most commonly supported (fig. 3B). It is noteworthy that the parallelism level among Mesoamerican highland populations was much stronger than observed between the Andes and other highland populations (fig. 3B). Not only do fewer loci show evidence of selection in pairs of populations including the Andes, but repeatedly selected loci are more likely to have arisen independently. The mean value of an independent source was 6.2% in Andean comparisons, in contrast to a mean of 2.0% within Mesoamerican highland population comparisons (fig. 3B). Previous studies have suggested that migration between the Andes and Mesoamerican populations is unlikely [36]. We therefore further evaluated all repeatedly selected SNPs by calculating the divergence statistic, *F_ST_*, between pairwise highland populations to assess how this varies based on the assigned model. Repeatedly selected SNPs consistent with a migration model exhibited weaker divergence (lower *F_ST_*; *p* ≈ 0; supplementary fig. S5, Supplementary Material online), an additional indicator of gene flow. Moreover, we calculated the 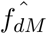 statistic (Malinsky et al. 2019) in 10-*kb* non-overlapping windows across the genome to further assess the signal of gene flow between the Andes and Mesoamerican highland populations. In a (((P1,P2)P3),O) shaped phylogenetic tree, the 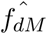 statistic gives positive values for introgression between P3 and P2 and negative values for introgression between P3 and P1. We found SNPs consistent with the migration model demonstrated more positive 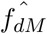 values, providing additional support for gene flow between the Andes and Mesoamerican highland populations (supplementary fig. S6, Supplementary Material online). In summary, while the proportion of dually selected SNPs between the Andes and Mesoamerican highland populations (from 7.8% to 11.2%) was consistent with previous work (7% - 8%) [36], the two most supported source models were standing variation and migration, suggesting a more prominent role for migration than previously hypothesized [36].

### Molecular parallelism at the genic level

We next evaluated the extent of molecular parallelism at the gene level. A gene was classified as an outlier if it contained *PBE* outlier SNPs within the gene or 10 kb upstream or downstream (supplementary table S3, Supplementary Material online). 1651 candidate genes were observed in the Mexican Highlands, among which 360 (21.8%) contained or were nearby outlier SNPs which were shared with at least one other highland population. The percentage dropped to 18.9% in the Guatemalan Highlands, 10.0% in the Southwestern US Highlands and 9.4% in the Andes (supplementary table S4, Supplementary Material online). We utilized the R package SuperExactTest [39] to evaluate the two- to four-degree intersection of outlier genes among highland populations. As observed in results at the SNP level, significant overlap was observed for each comparison (supplementary fig. S7, supplementary fig. S8, Supplementary Material online). In many instances, common outlier genes involved selection on different SNPs across highland populations. For example, 349 (21.1%) of the 1651 selected genes in the Mexican Highlands were in common with other highland populations, but showed evidence of selection on distinct SNPs. The percentage was 18.8% in Guatemala, 24.7% in the Southwestern US, and 18.5% in the Andes (supplementary table S4, Supplementary Material online).

#### Constraint of the adaptation target size leads to molecular parallelism

There are two explanations for the extent of molecular parallelism we have observed in genes selected in the highlands [9]. First, it is possible that adaptation is constrained by the number of genes that can affect a trait, effectively placing physiological limits on the routes adaptation can take. Alternatively, many different genes may have the potential to affect the trait, but deleterious pleiotropic effects of variation in many genes may prevent them from playing a role in adaptation. We will refer to these possibilities as “physiological” and “pleiotropic” constraint. Consistent with our analysis above, Yeaman [9]’s *C*_χ^2^_ statistic finds strong evidence that selection at the same loci is shared among populations more often than can be explained by chance (*C*_−χ^2^_ from 19.6 to 40.1 with a mean of 28.05, permutation *p* < 0.0001). To evaluate the role of physiological constraint in parallel adaptation, we tested an alternative null model which assumes the set of genes selected in at least one population represents all genes that can possibly affect the trait. We then tested whether there was still evidence of excessive sharing after accounting for physiological constraint [9]. We found *C*_χ^2^_ was less than 0 in all highland pairs except the Mexican and Guatemalan Highlands pair (*C*_χ^2^_ = 1.9, *p* = 0.025). These results indicate that repeated selection of genes among most highland pairs may simply be due to physiological constraint in the number of loci that can contribute to variation in a trait. To explore this possibility further, we assessed the prevalence of molecular parallelism in the flowering time pathway, a well-characterized gene network known to be of adaptive importance in highland environments.

### Molecular parallelism within the flowering time pathway

Flowering time in maize is known to be a highly quantitative trait and an example of polygenic adaptation [40, 41, 42, 43]. As such, we may expect that highland adaptation for flowering is the result of subtle, coordinated allele frequency shifts across many loci. In evaluating flowering time genes we tested: 1) the strength of selection on loci associated with flowering time relative to expectations based on drift, and 2) the intersection of selected genes in the pathway in multiple highland regions. To first identify whether flowering time was targeted during highland adaptation, we conducted a genome-wide association study (GWAS) of 29 traits measured in short-day conditions in Ponce, Puerto Rico and Homestead, Florida [44] using a maize association mapping panel comprised of tropical and temperate inbred lines [45]. We then summarized the frequency of multiple alleles affecting a trait of interest with a polygenic score and tested for associations between polygenic score and elevation of origin of our landraces, while controlling for variation in relatedness between samples following Josephs et al. [43]. A significant result implies that allele frequencies that increase trait value are more common at one end of the elevation gradient (either high or low) than expected due to drift. Seven traits showed a significant relationship between their conditional polygenic score and elevation with a false discovery rate < 0.1. These traits were primarily related to flowering time: Days to Silk, Growing Degree Days to Silk, and Ear Height from ’06 Puerto Rico, Growing Degree Days to Silk, Growing Degree Days to Tassel, and Ear Height in ’06 Florida, and Growing Degree Days to Silk in ’07 Florida (fig. 4).

**Figure 4:**
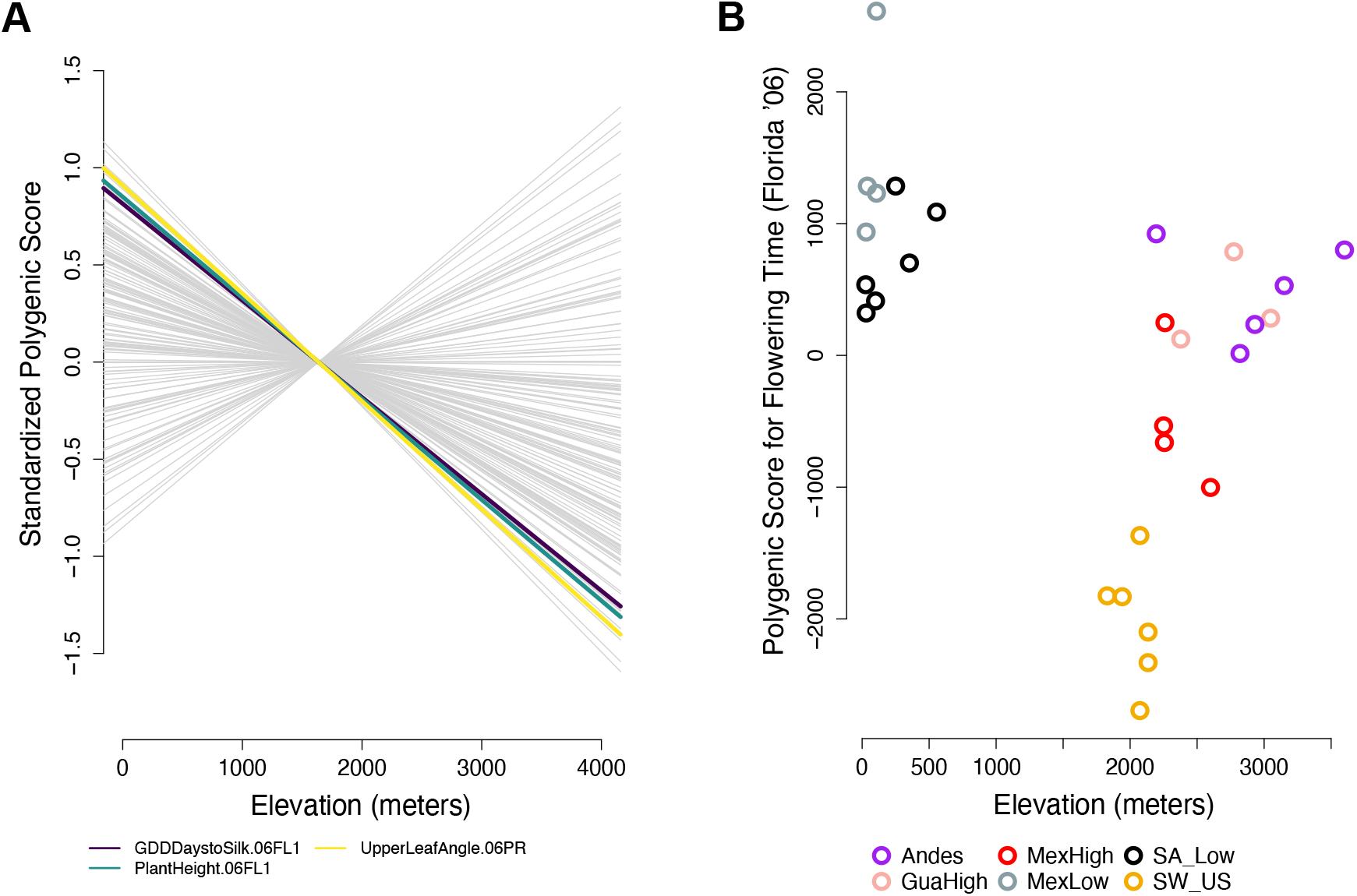
Polygenic adaptation along elevation in the landraces. A. A linear regression between polygenic score for all 29 trait-environment combinations tested and elevation of origin. The lines for the seven traits that showed significant signals of polygenic adaptation are colored and all other traits are shown in gray. B. Polygenic score for days to silk in Florida (2006) for all landraces is negatively correlated with elevation of origin.

Given this support for selection on flowering time during highland adaptation, we explored the overlap between our selected genes and known flowering time candidates. First, we utilized a list of 904 maize flowering time candidate genes aggregated by Li et al. [46] (supplementary table S5, Supplementary Material online). Within this list, we found a substantial excess of flowering time genes targeted by selection among groups of two, three, and four highland populations (p < 2.87*e* – 5, hypergeometric test; fig. 5A, supplementary table S6, Supplementary Material online). Outlier SNPs within these genes also showed strong co-directional changes in allele frequency across highland populations (91.8% – 100%) (supplementary fig. S9, Supplementary Material online). As observed in our genome-wide analysis, candidate sharing at flowering time loci was stronger among Mesoamerican highland populations than between comparisons with the Andes (fig. 5). We also applied the *C*_χ^2^_ statistic to this list of genes and confirmed enriched sharing of selected genes among highland populations (mean *C*_χ^2^_ 6.618, permutation p < 0.0001). Our finding of an excess of shared candidates across regions within a large set of known genes that affect flowering time, suggests physiological constraint alone cannot explain patterns of convergence, but instead that variation at only a subset of genes can affect flowering without deleterious pleiotropic consequences.

**Figure 5:**
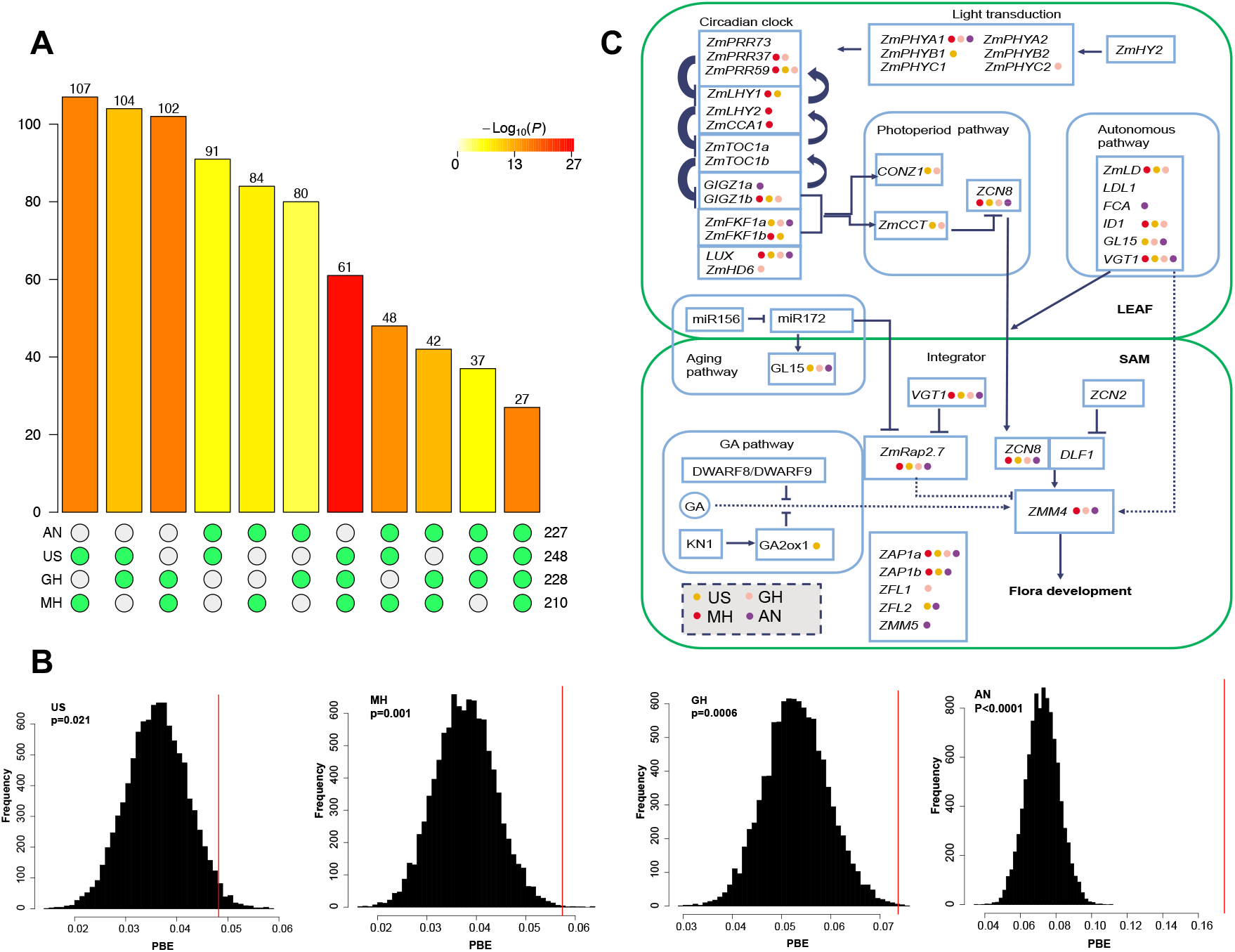
Convergent adaptation in the flowering time pathway. A. Intersection of flowering time outlier genes among four highland populations. B. Distribution of mean *PBE* values of SNPs located within and in the 10-kb flanking regions of core flowering time genes [47] (the red line) against genomic background (the black bars). C. Diagram showing selected genes in the flowering time network. Colored dots indicate the population(s) in which selection was detected.

We then narrowed our focus to a subset of core flowering time genes [47] with highly characterized functional roles (fig. 5C). Thirty-two of the 39 core flowering time genes were selected in at least one highland population, which indicated over-representation of flowering time genes among all selected genes (p = 0.005, hypergeometric test). While common outlier genes between the Andes and other highland populations were not significantly enriched in this smaller list (*p* > 0.074, hypergeometric test; supplementary table S6, Supplementary Material online), enrichment was observed among all other two and three highland-population comparisons (*p* < 0.042, hypergeometric test; supplementary table S6, Supplementary Material online). These results were confirmed through calculation of the *C*_χ^2^_ statistic (Mesoamerican highland population comparisons, mean *C*_χ^2^_ 2.32, permutation p < 0.01; comparisons to the Andes, mean *C*_χ^2^_ 1.575, permutation p from 0.015 to 0.054).

The *PBE* values of outlier SNPs within or in the 10-kb flanking regions of flowering time genes were in the extreme tail of the genome-wide distribution (fig. 5B), with some of the best-characterized flowering time genes in maize clearly targeted by selection in multiple highland populations (fig. 5C). For example, ZCN8 (GRMZM2G179264), which is orthologous to FLOWERING LOCUS T (FT) in *Arabidopsis*, has been shown to play a significant role in maize adaptation to high latitudes [48] and was selected in all four highland populations in our study. Floral transition is regulated by several MADS-box genes, particularly ZMM4 (GRMZM2G032339), which is under selection in all but the Southwestern US Highlands. VGT1 (GRMZM2G700665; vegetative to generative transition 1) was identified as an outlier in all four highland populations, with the same non-synonymous SNP (8:131578990, indicating locus 131578990 on chromosome 8) targeted in the Mexican and Guatemalan Highlands (supplementary fig. S10, Supplementary Material online). In contrast, in the Southwestern US Highlands, the selected SNP (8:131580179) in VGT1 was located in the 3’ UTR region and, in the Andes, the selected SNP (8:131579463) was a synonymous mutation near the 3’ UTR (supplementary fig. S10A,B,C, Supplementary Material online). Two CONSTANS (CO) genes, CONZ1 and ZmCCT, were selected in both the Mexican and Guatemalan Highland populations. We further found Gigantea2 (GRMZM5G844173) was selected in all but the Guatemalan Highlands and FKF2 (GRMZM2G106363) in all but the Mexican Highlands. Among circadian clock genes, LUX (GRMZM2G067702)(supplementary fig. S10D, Supplementary Material online) was selected in all four highland populations and ZmPRR59 was an outlier in the three Mesoamerican highland populations. In addition, the zmZTLa - F box protein ZEITLUPE (GRMZM2G113244), RVE2 (GRMZM2G145041) and PRR5 (GRMZM2G179024) were selected in all four highland populations. Finally, the light transduction gene PHYA1 (GRMZM2G157727) was targeted by selection in all but the Southwestern US highland populations.

## Discussion

The prevalence of parallel evolution is a long-standing question in the field of evolution. Through a comprehensive evaluation of parallel adaptation among independent maize highland populations, we found that, while most adaptation is independent, parallelism is more common than expected. We observed a particularly high level of parallelism among Mesoamerican highland populations relative to comparisons including Andean maize. The vast majority of parallel adaptive alleles we discovered have risen in frequency in the highlands from migration and standing variation, and only a small percentage represent *de-novo* mutations. However, the proportion of repeatedly selected SNPs from *de-novo* mutations is highest in pairs of populations that include the Andes. Our analyses further reveal that adaptive routes from genotype to phenotype have been canalized by both physiological (*i.e*., few genes contributing to a trait) and pleiotropic (*i.e*., a subset of causal genes can contribute to trait variation without negative pleiotropic consequences) processes. For example, we observe that the flowering time pathway is saturated with selected genes potentially contributing to highland adaptation, with many of these loci showing evidence of selection in multiple highland regions, particularly in Mesoamerica.

Approximately one quarter (mean 24.0%) of highland adaptation SNPs showed a pattern of molecular parallelism in Mesoamerica. A smaller percentage also showed this pattern in Andean comparisons (mean 10.7%), but this represents a slightly higher proportion than previously reported (7% - 8% between the Andes and the Mexican Highlands) [36]. We have also expanded upon previous studies of parallel highland maize adaptation [36] in a number of ways. Our study has broadened the geographic scope of sampling for highland maize populations and is the first such study using high-depth, whole-genome resequencing data, which allowed us to attain a more complete picture of parallelism. Additionally, our use of the PBE statistic allowed for detection of selection specifically in the highlands and reduced the extent of false positives, a limitation of studies that focus solely on population differentiation outliers. Our polygenic adaptation analyses also helped to clarify specific phenotypes targeted by selection during adaptation, showing clear evidence of selection on flowering time. We have also uncovered a pervasive role of migration in transferring adaptive alleles across highland regions. In fact, our analyses revealed a moderate proportion of parallel adaptation through migration even between the Andes and other highland populations, in contrast to previous studies that appeared to exclude this possibility [36] based on simulations. This finding is consistent with a recent study which showed that some traits, for example flowering time, may not demonstrate a GxE effect [42], allowing adaptive alleles to move between regions through a matrix of habitat (*e.g*., the lowlands) where they do not clearly confer an adaptive advantage.

The extensive parallelism in maize highland adaptation has likely been affected by multiple factors and is consistent with expectations based on population genetic theory. First, a high level of genetic diversity in the species provides substantial standing genetic variation and potentially beneficial mutations. Similarly, Zhao et al. [49] showed that the more genetically diverse species *Drosophila hydei* contained more adaptive alleles than *D. melanogaster*. Second, the recent timing of expansion and divergence in maize has likely affected the extent of parallelism. The probability of parallelism will decrease when populations diverge over a greater period of time, as less shared standing variation would be expected to be the source of adaptation. Increased divergence time would also allow for novel beneficial mutations to arise, potentially in distinct genes and pathways [50]. For example, Preite et al.[51] demonstrated a moderate level of molecular parallelism in adaptation to calamine metalliferous soils within *Arabidopsis* species, but less parallelism between species. In addition, a recent study [52] discovered that the degree of molecular parallelism decreases with increasing divergence between lineages in *Arabidopsis* alpine species. Third, population genetic modeling predicts that traits with pleiotropic constraint are more likely to demonstrate molecular parallelism [53]. Despite the quantitative nature of flowering time in maize, the *C*_χ^2^_ statistic still showed evidence of genetic constraint and convergence even after limiting our null to genes in the known flowering time pathway. The genetic parallelism could also be attributed to the high effective population size of the species and extensive migration between highland regions. Similarly, a recent study found molecular parallelism for cold tolerance between distantly related species (lodgepole pine and interior spruce), indicating certain key genes playing crucial roles in cold adaptation [54]. Finally, theory predicts that gene flow may have contrasting impacts on parallel adaptation. Gene flow between differentially adapted populations may introduce maladaptive alleles, counteracting the effects of local selection [53]. For example, Holliday et al. [55] found islands of divergence in *Populus trichocarpa* populations across altitudinal clines and proposed that coadapted genes in strong linkage may buffer against genetic introgression from maladapted individuals. Our results, consistent with earlier findings for maize in the highlands of Mexico [36, 37], argue instead for an important role for adaptive introgression in increasing the likelihood of parallelism. Similarly, adaptive introgression at the EPAS1 locus has been shown to underlie convergent highland adaptation in both humans [56] and dogs [57].

In summary, the emerging story of parallel highland adaptation in maize is consistent with existing theory. Gene flow is common across maize populations, but does not appear to swamp locally adaptive alleles. Rather, highland adaptation appears to have initially been facilitated in the Central Plateau of Mexico by introgression from the locally adapted wild species, *mexicana* [37]. Subsequently, gene flow from the Mexican Highlands appears to have contributed to highland adaptation in other regions of the Americas [38], consistent with our finding of migration as the major source of molecular parallelism. Novel alleles from *mexicana* augmented the extensive standing variation already found within the species [26], providing a pool of adaptive variants that appear to have independently risen in frequency in multiple highland regions.

Similar adaptive processes (*e.g*., gene flow with newly encountered, locally adapted populations and species, repeated selection on standing variation in independent but similar habitats) likely occurred as other crops expanded from their centers of origin [58]. However, crops that were domesticated from wild species with small effective population sizes, with more pronounced domestication bottlenecks, or without widespread, locally adapted congeners may have lacked the adaptive potential necessary to achieve the broad and varied distribution of maize. The success of invasive species has also been linked to hybridization [59] and high levels of standing variation [60]. A thorough understanding of adaptive processes during rapid expansion across diverse habitats may therefore inform invasive species mitigation strategies, assisted migration in the face of climate change, and crop breeding for tolerance of extreme environmental conditions. Highland adaptation in maize is a clear example in which adaptation has drawn substantially from a shared pool of variants, with the same alleles, genes, and pathways contributing to the evolutionary success of the species across continents and millennia.

## Materials and Methods

### Samples and Data

Samples and SNPs were from our previous publication [38]. SNPs were removed when located in a known inversion, large introgression regions from *mexicana* and centromeric regions, and later filtered to retain only those with sequencing depth greater than 10X and minor allele frequency greater than 0.05 (supplementary fig. S11, Supplementary Material online). Nineteen environmental variables were obtained from the WorldClim data set (http://www.worldclim.org/) and were utilized to perform a Principal Component Analysis (PCA) with the vegan package in R [61]. We employed Variant Effect Predictor (VEP) from Ensembl [62] to further annotate genomic location and functional consequences of SNPs.

### *PBS* and *PBE* calculation

Compared to the commonly used statistic to estimate levels of population differentiation - *F_ST_*, *PBS* calculates allele frequency change specific to one focal population by adding pairwise *F_ST_* values involving a third “outgroup” population [6].

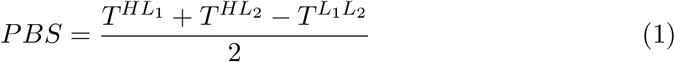

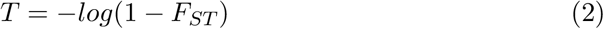

*PBE* is a recently described derivative of *PBS*, which overcomes the limitation of *PBS* values being high when all populations have long branches. *PBE* instead quantifies the difference of the observed and the expected *PBS* values [63].

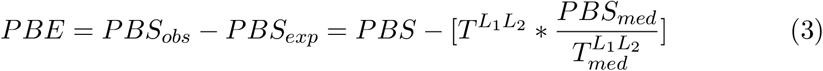

Here, *T*^*L*_1_*L*_2_^ quantifies genetic differentiation of the two non-focal populations. *PBE* is expected to be strongly positive when selection acts specifically on the focal population.

We calculated *PBE* for filtered SNPs among one highland population, its corresponding lowland population (the Mexican Lowland population was paired with the Mesoamerican Highland populations; the South American Lowland population was paired with the Andes population) and the *parviglumis* Palmar Chico population (supplementary fig. S11, Supplementary Material online).

Pairwise *F_ST_* values were calculated in vcfTools [64] and custom R scripts were used to compute *PBS* and *PBE* values of the focal highland population. SNPs with *PBE* values higher than the 95% quantile of its distribution were regarded as outliers and qualitative results were confirmed in the 99% quantile. The R package “SuperExactTest” [39] was utilized to evaluate if the overlap of outlier SNPs between pairwise highland populations was enriched.

In addition, we also evaluated if the same allele is elevated to high frequency among outlier SNPs between pairwise highland populations (supplementary fig. S12, Supplementary Material online). If there is no allele frequency change between the highland and its lowland counterpart, we categorized the SNPs as “non-directional”. When a SNP was non-directional in both highland populations being compared, it was removed from the analyses. When the same allele was at elevated frequency in both highland populations compared to their lowland counterparts, the SNP was categorized as “co-directional”. When different alleles were at high frequency across two highland populations, the SNP was categorized as “anti-directional”. We then compared the proportion of co-directional SNPs between the outlier and neutral SNPs.

In order to exclude the bias of genetic drift on the directionality of changes in allele frequency, we approximated the two-dimensional site frequency spectrum (2dsfs) of outlier SNPs using a subset of the neutral SNP set. We divided the nonreference allele frequency in Mexican Lowland maize and *parviglumis* into ten equal bins in both the outlier and neutral SNP set. Then we matched neutral SNPs with the same ancestral 2dsfs (representing the combination of allele frequencies in both the Mexican Lowland and *parviglumis* populations) with the outlier SNP set and sampled the same amount of SNPs in each allele frequency bin. Last, we tabulated the directionality of the randomly sampled neutral SNPs to check how ancestral allele frequency influenced the proportion of co-directional SNPs.

### Composite likelihood calculations to distinguish the mode of parallelism

Following the method developed and applied in [11], we calculated the composite likelihood under four models for the outlier regions shared among all four highland populations, after thinning for linkage. The four models tested were as follows: 1) neutrality (no selection), 2) independent mutations of the beneficial allele in four highland populations, 3) a single origin of the beneficial allele in one single highland population and spread via gene flow into the other population during the sweep, 4) standing variation present in the ancestor of the four highland populations and spread via gene flow into other populations some time *t* before selection occurred. The details of these four models can be found in [11]. For each outlier, we obtained SNPs in the surrounding 20-kb window and calculated the composite likelihood under each of the four models for this dataset. We compared composite likelihoods under the models for each outlier and obtained maximum composite-likelihood-parameter estimates, specifically for the standing time *t* in the fourth model, to assess the timing of gene flow. Similarly, we also evaluated the composite likelihood values under four models for each set of dually selected SNPs in pairs of highland populations.

### Polygenic adaptation for quantitative traits

We followed the methods developed by Berg et al. [65] and Josephs et al. [66] to detect polygenic adaptation in the highland populations. Briefly, these methods detect adaptive divergence for a trait by 1) finding loci associated with that trait in a GWAS, 2) summarizing the allele frequencies and effect sizes at these loci in the populations of interest using a polygenic score, and 3) testing to see if the association between these polygenic scores and an environmental character (in this case, elevation) are greater than could be explained by drift.

To identify loci associated with traits that could be under selection, we conducted genome-wide association studies using a maize panel developed for GWAS, often referred to as “the 282” or “the Major Goodman panel” [45]. Single nucleotide polymorphisms (SNPs) for 263 individuals in the Major Goodman panel were called from whole genome sequencing data from Bukowski et al. [67], removing individuals with genotype calls for <70% of polymorphic sites. Of all the SNPs called in Bukowski et al.[67], we only used those that were also polymorphic in the maize landraces and that had a MAF > 0.01 and were missing data for <5 % of individuals, leaving us with approximately 5 million SNPs per test. We used trait measurements made in three short-day common garden experiments, from Florida in 2006 and 2007 and Puerto Rico in 2006, from Hung et al. [44] where there were data available for at least 80% of the 263 individuals, leaving us with 29 trait-environment combinations. We conducted GWAS using GEMMA with default parameters [68], with a standardized kinship matrix to control for population structure.

We used the GWAS hits to generate polygenic scores for each individual in the landrace panel using all loci with a p value < 0.1 that had been pruned down to the strongest hit per 1 cM window using a linkage map from Ogut et al. [69], leaving us with an average of 216 SNPs per trait (range = 147-289). If there are *M* loci associated with a trait, each with effect size *α_m_*, *p_im_* is the allele frequency of the *m^th^* allele in line *i*, which will either be 0, 0.5, or 1, then we can calculate the polygenic score for the *i^th^* line as *Z_i_* where

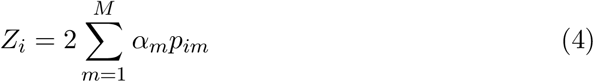

Shared population structure between the GWAS panel and the test panel can lead to false positive signals of selection [66]. To test for shared structure between the GWAS panel used here and the landraces, we constructed a joint kinship matrix between all samples in both panels, following the same methods used in Josephs et al. [66]. Plotting the first two eigenvectors of this joint kinship matrix (the principal components) show that there is likely shared population structure between the landraces and the GWAS panel (supplementary fig. S13A, Supplementary Material online). We constructed a conditional kinship matrix that estimates the amount of relatedness between landraces conditional on relatedness captured in the GWAS panel, and found that this new conditional matrix explains less variation than the original matrix made from landraces alone (supplementary fig. S13B, Supplementary Material online). This finding is further evidence that there is shared population structure between the two panels.

In light of this shared structure, we conducted a conditional test, following Josephs et al. [66]. Let 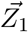 be the vector of *Z_i_* for the 31 highland and lowland landraces discussed in this paper and 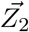 is the vector of *Z_i_* for the individuals used in the GWAS. We model the combined vector of polygenic scores in both panels as

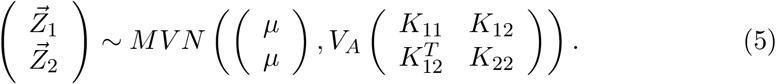

where, *μ* is the mean of the combined vector [*X*_1_, *X*_2_], *K*_11_ and *K*_22_ are the kinship matrices of the genotyping and GWAS panels, and *K*_12_ is the set of relatedness coefficients between lines in the genotyping panel (rows) and GWAS panel (columns).

The conditional multivariate null model for our polygenic scores in the landraces conditional on the GWAS panel is then

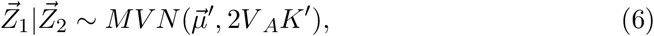

where 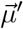 is a vector of conditional means with an entry for each sample in the genotyping panel:

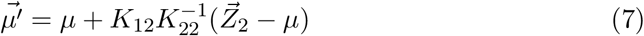

and *K*′ is the relatedness matrix for the genotyping panel conditional on the matrix of the GWAS panel,

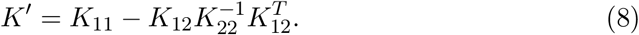

To test for selection, we calculate the difference between observed polygenic scores and conditional means as 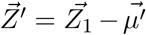. Higher values of 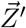 indicate that the observed polygenic score in a landrace individual is greater than would be expected based on relatedness between that landrace individual and individuals in the GWAS panel.

After calculating 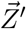 for 29 environment-trait combinations, we test for a correlation between 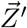 and the elevation of origin of each landrace beyond what would be expected due to neutral drift using the methods described in [65]. We transformed both 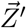 and the mean-centered vector of elevations for each landrace by the Cholesky decomposition of the kinship matrix (*C*) using the following equation (shown here for 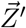)

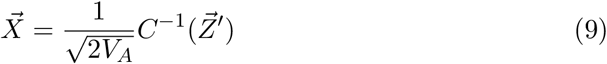

We estimate *V_A_* using the allele frequencies and effect sizes of the GWAS loci, using the following formula. If 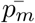 is the allele frequency of the *m^th^* locus across all individuals and *α_m_* is the effect size of that locus,

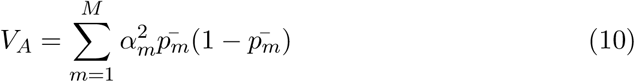

We then test for a linear relationship between 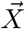 and the similarly transformed vector of elevations using the lm function in R [70] and control for the 29 tests done using Qvalue [71].

## Acknowledgments

This study was supported by Shenzhen Science and Technology Program (Grant No. KQTD2016113010482651), special funds for science technology innovation and industrial development of Shenzhen Dapeng New District (Grant Numbers RC201901-05 and PT201901-19), the US Department of Agriculture (USDA #2009-65300-05668), the National Science Foundation (NSF IOS #1546719; NSF IOS #1523733 to EB), USDA Hatch project (CA-D-PLS-2066-H), and startup funds from Iowa State University. We thank Graham Coop and members of the Hufford Lab for helpful comments and suggestions.

## Supporting Information

**Figure S1:**
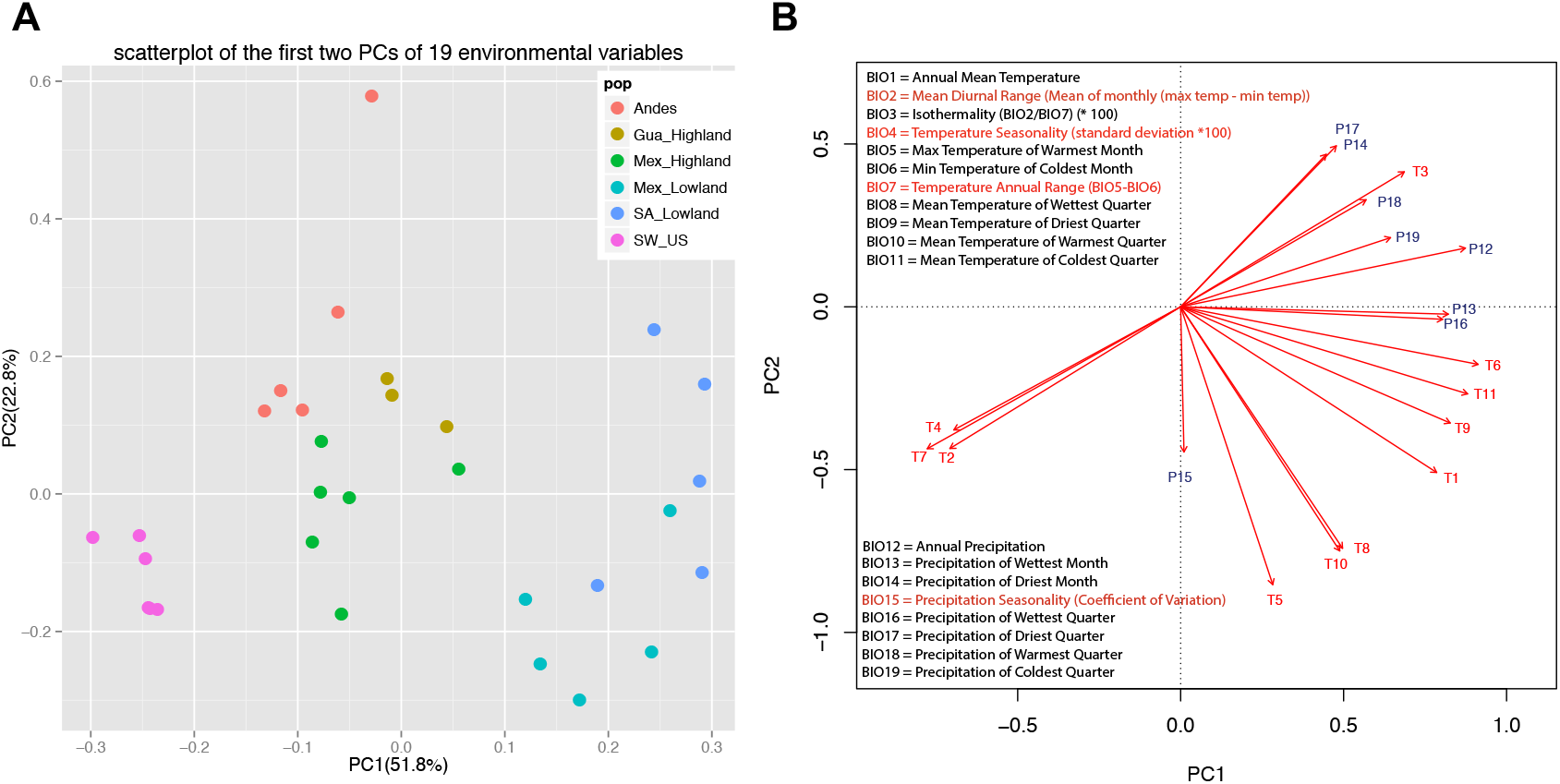
Environment of maize populations. A. PCA of 19 bioclim environmental variables for our samples. B. Projection of the 19 environmental variables on the first two PCs. Abbreviations for populations: GuaHigh, Guatemalan Highlands; MexHigh, Mexican Highlands; MexLow, Mexican Lowlands; SA_Low, South American Lowlands; SW_US, Southwestern US Highlands.

**Figure S2:**
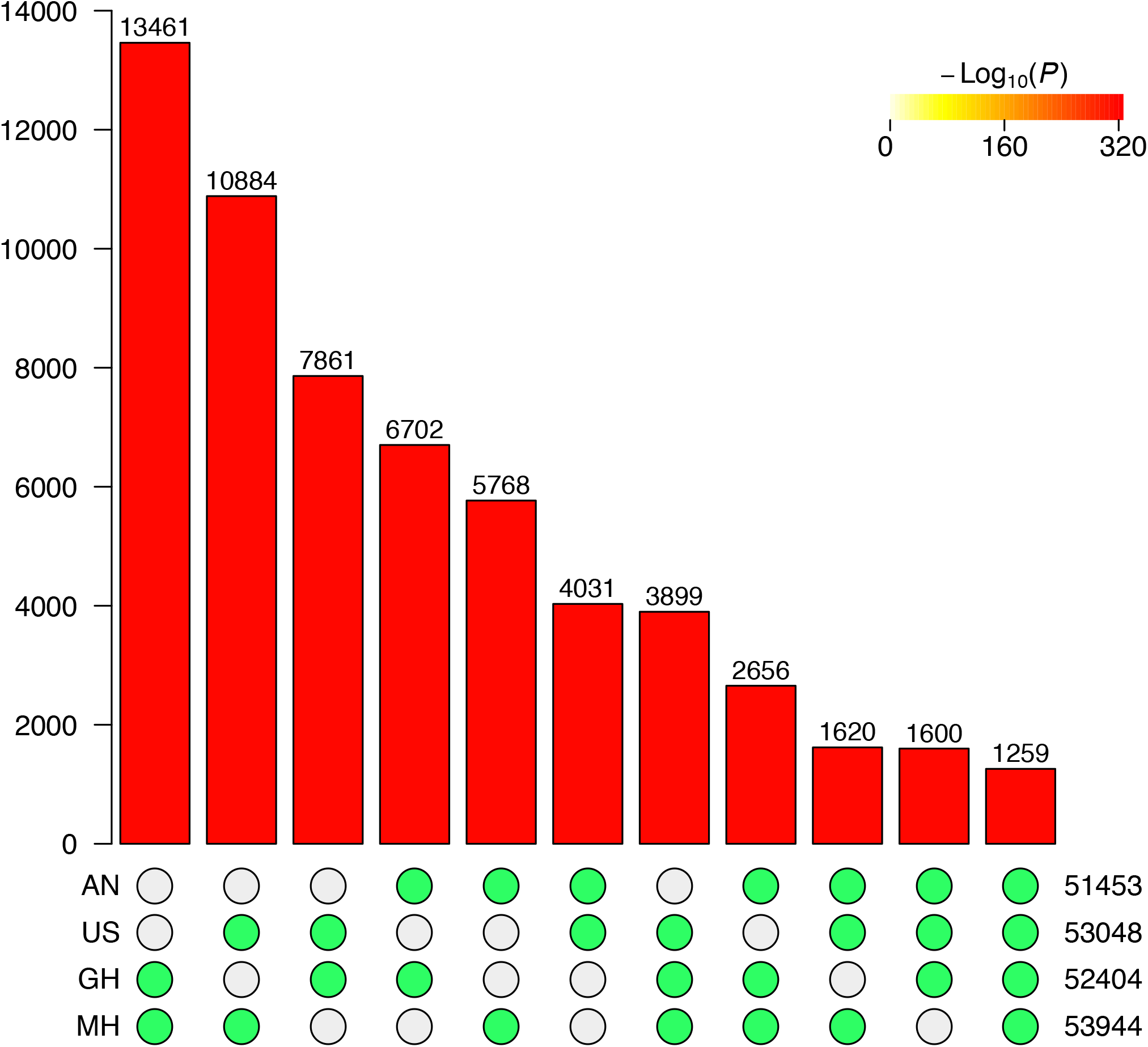
Intersection of outlier SNPs among four highland populations.

**Figure S3:**
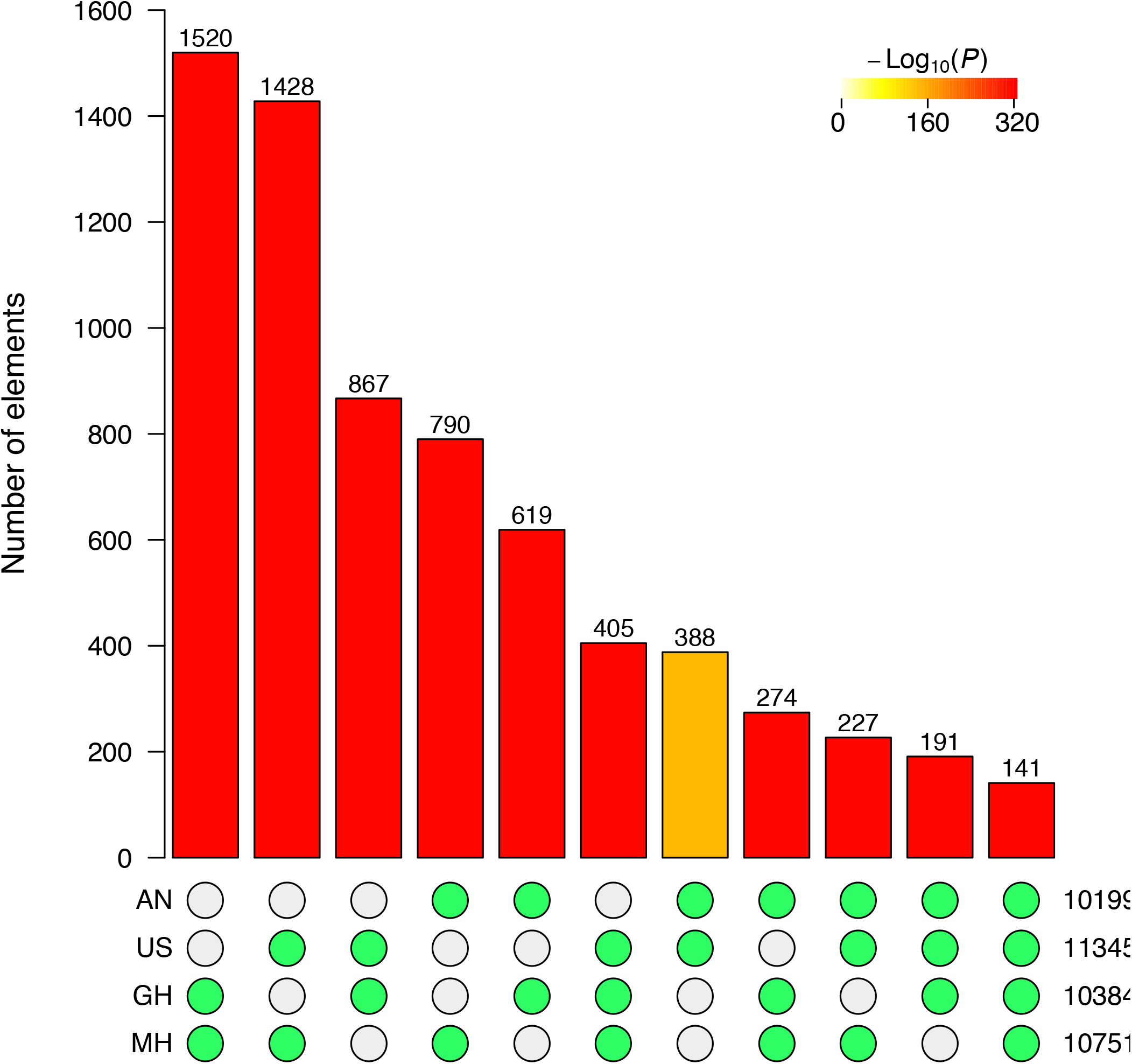
Intersection of outlier SNPs (1% top PBE values) among four highland populations.

**Figure S4:**
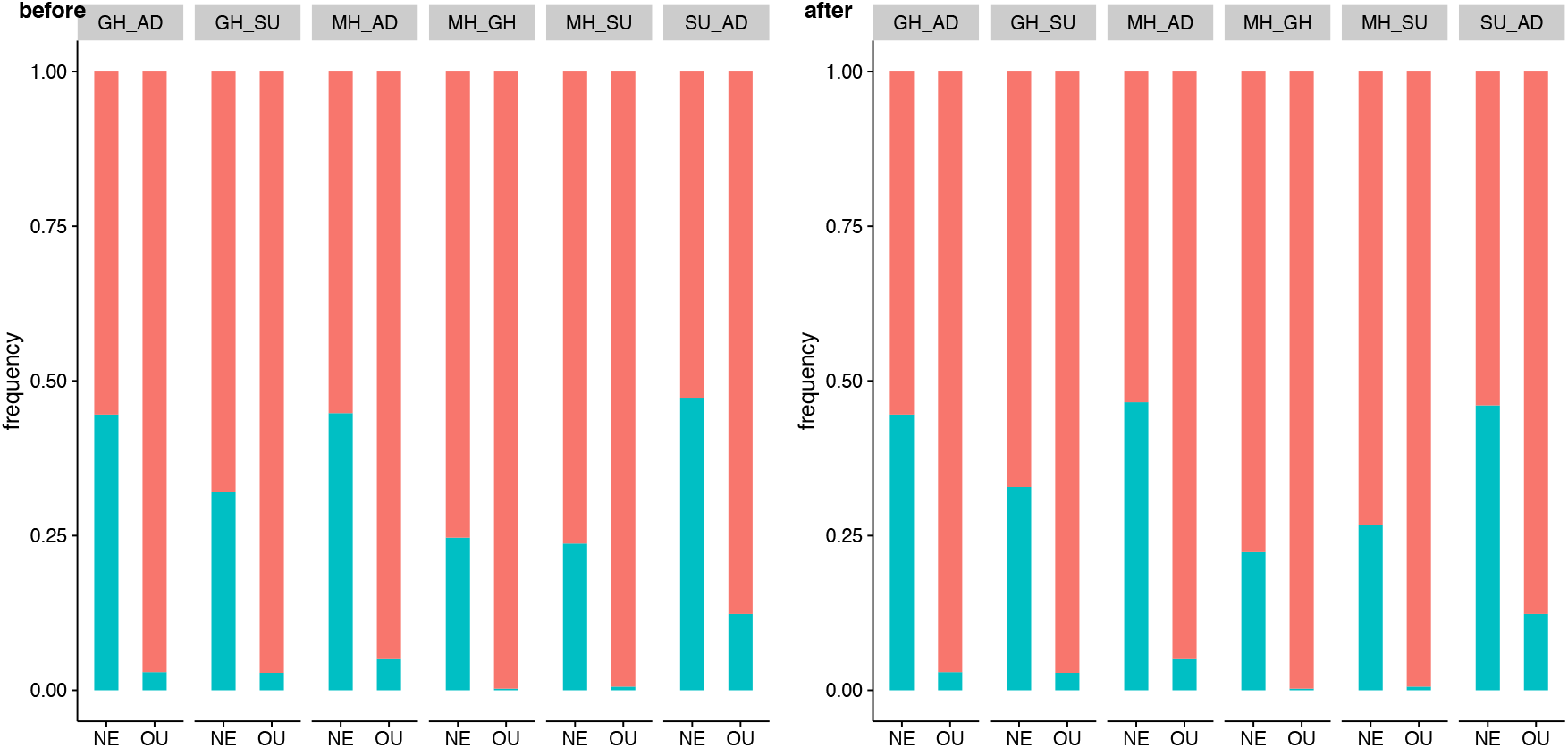
Ratio of co-directional and anti-directional SNPs in common neutral (NE) and outlier (OU) SNP set before and after correcting for 2dsfs.

**Figure S5:**
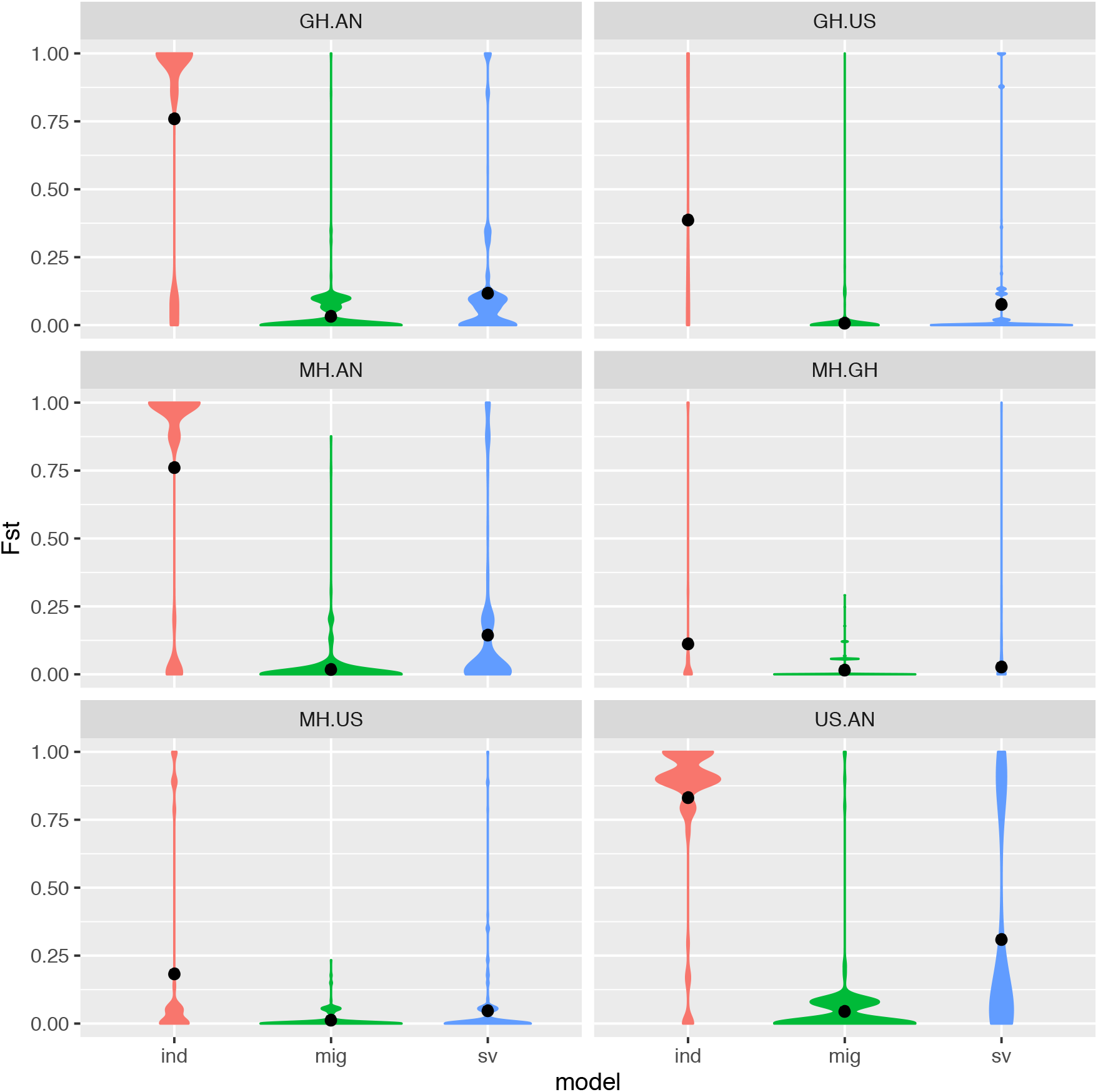
Comparison of Fst between pairs of highland populations for repeatedly selected SNPs assigned to different source models.

**Figure S6:**
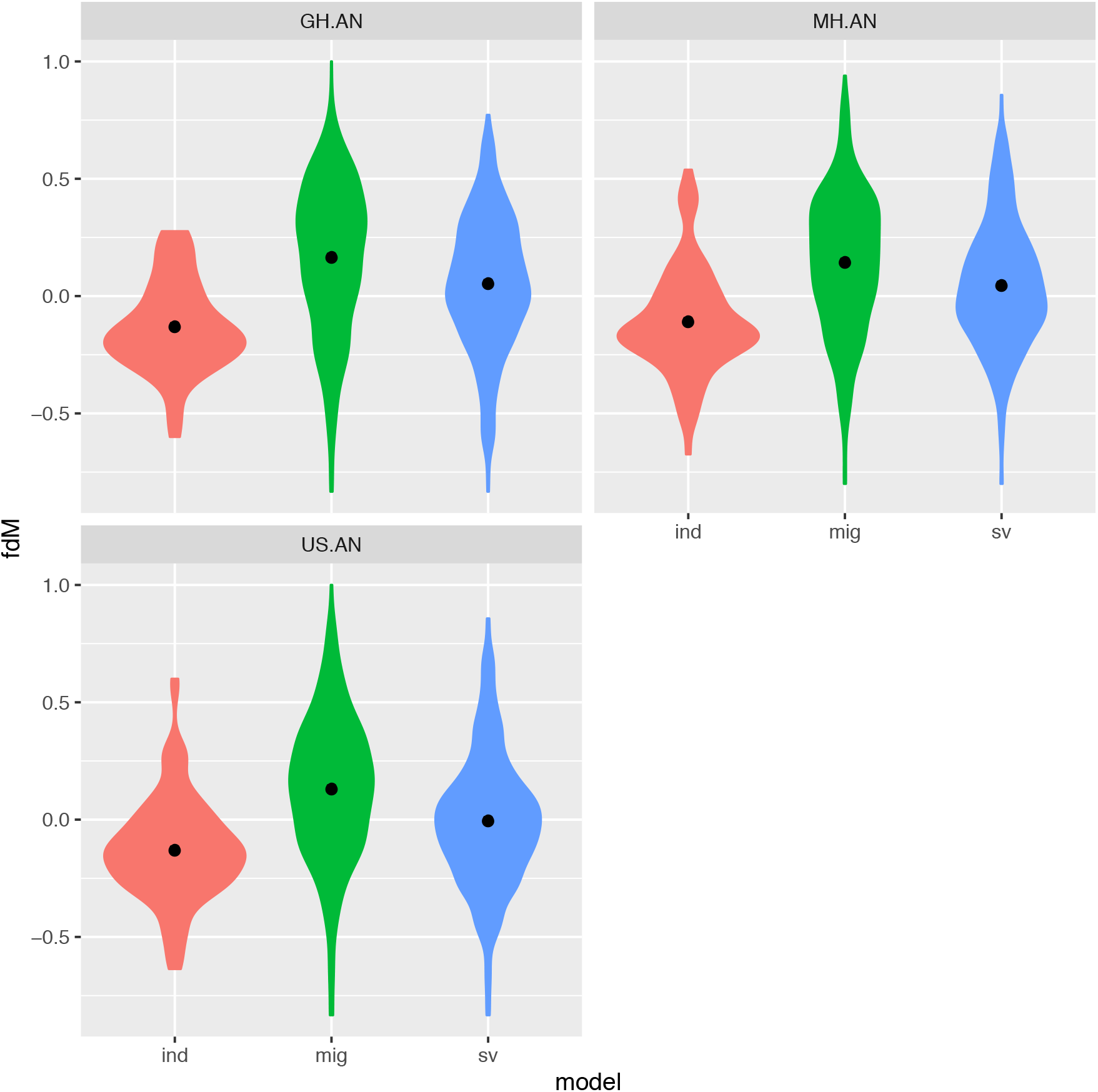
Comparison of 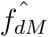 between pairs of highland populations for repeatedly selected SNPs assigned to different source models.

**Figure S7:**
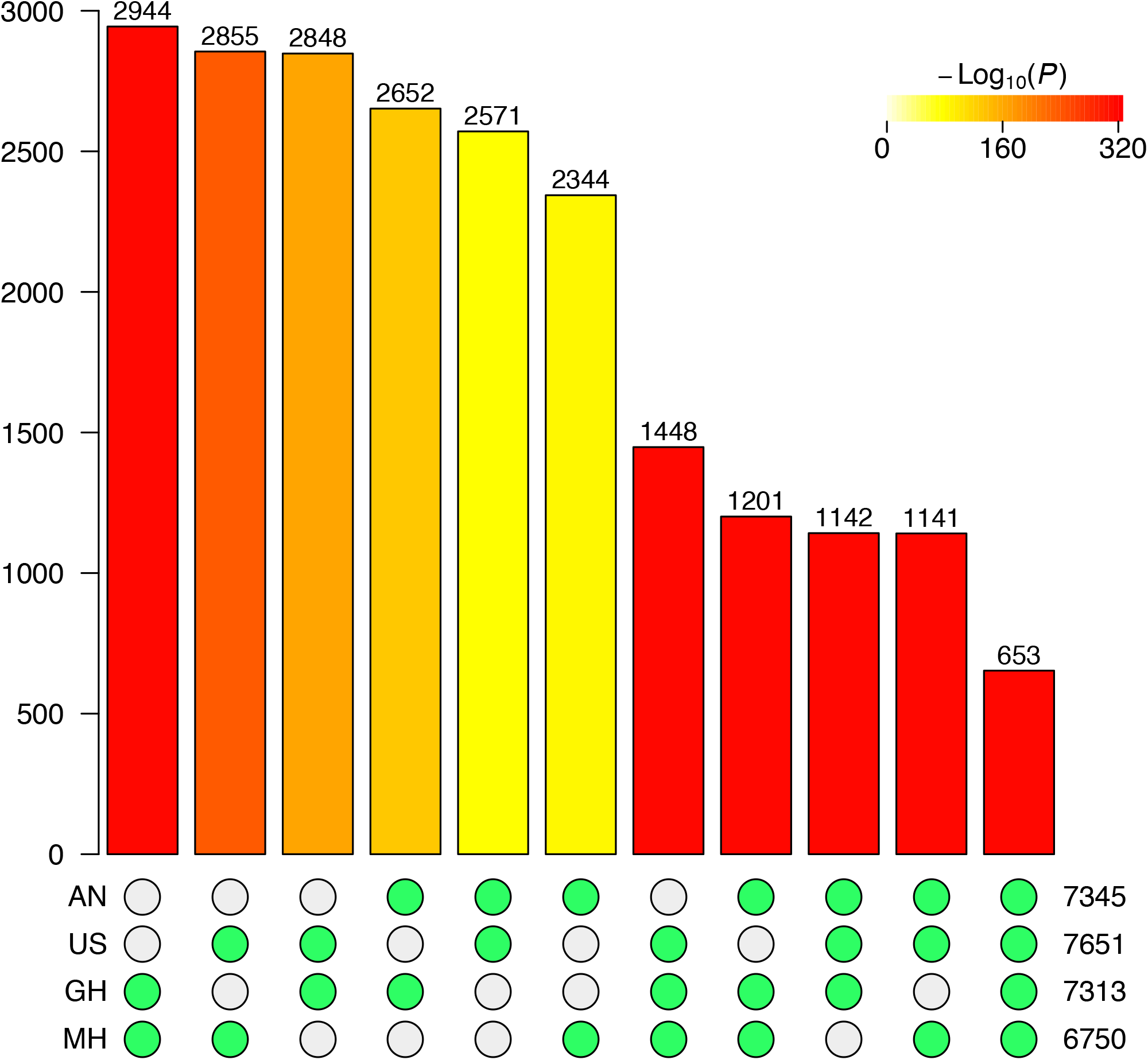
Intersection of outlier genes (based on SNPs with the top 5% PBE values) among four highland populations.

**Figure S8:**
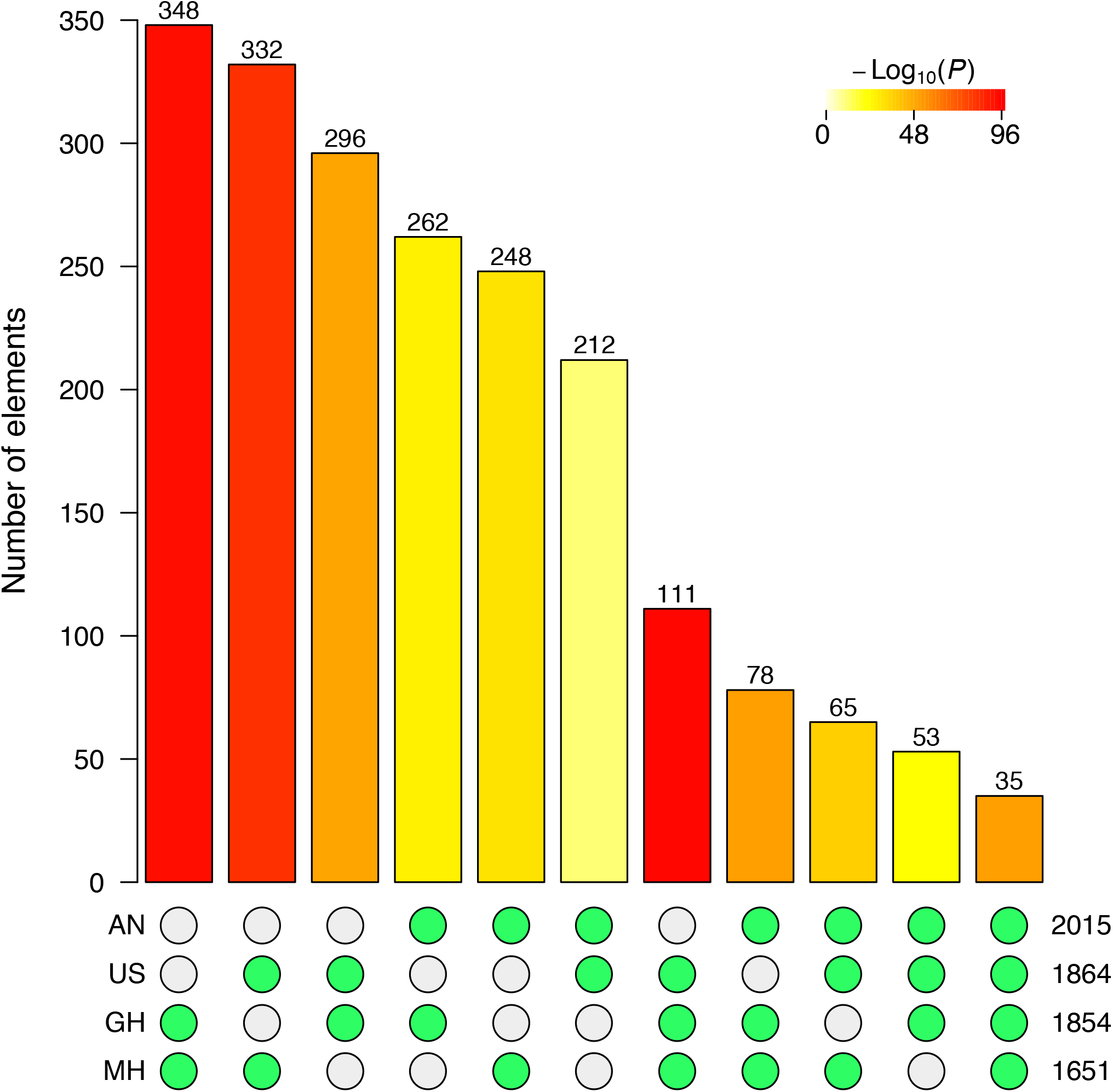
Intersection of outlier genes (based on SNPs with the top 1% PBE values) among four highland populations.

**Figure S9:**
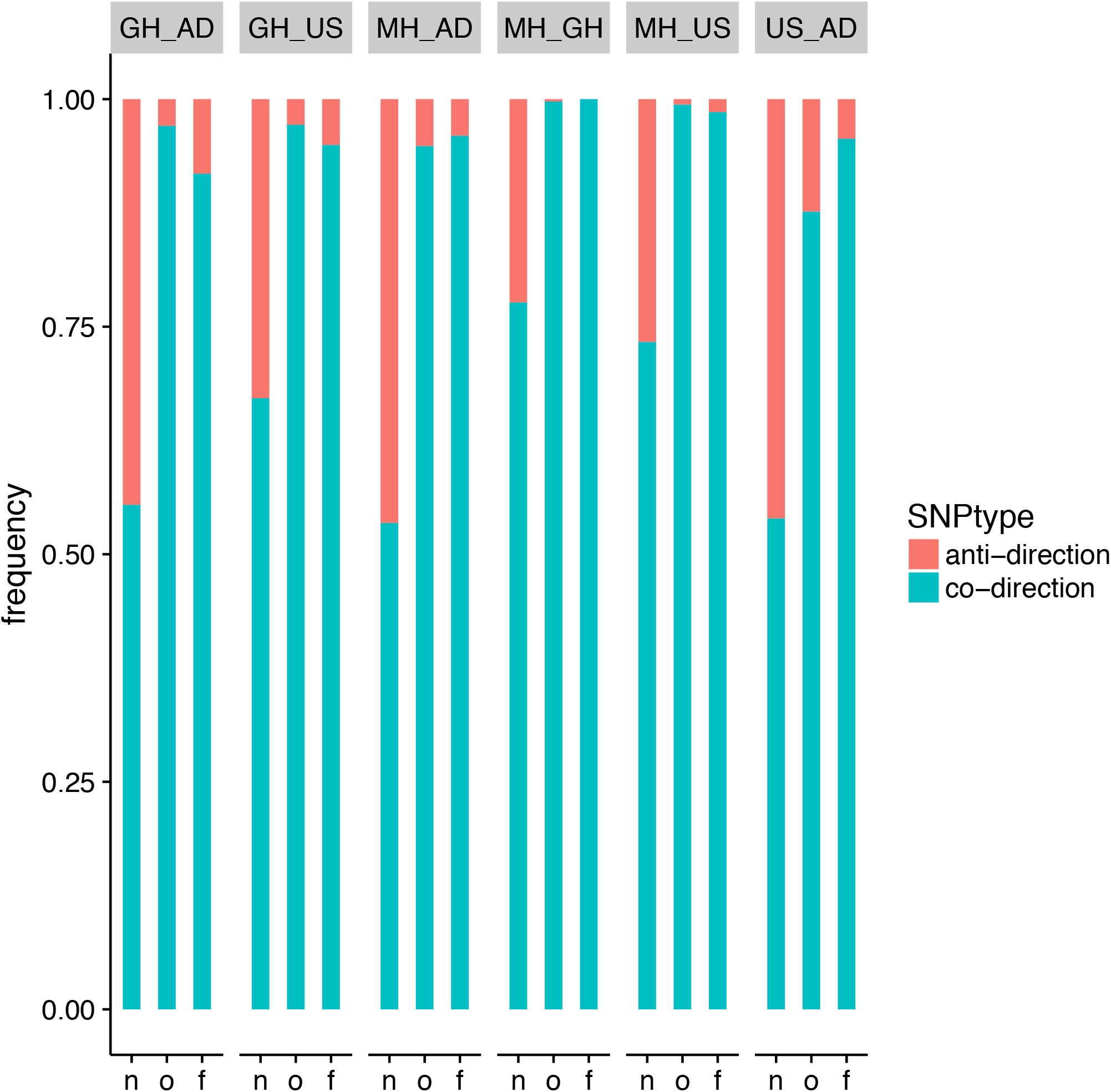
A barplot showing the dominance of co-directional SNPs in flowering time metabolism pathways. n: neutral SNPs; o: outlier SNPs; f: outlier SNPs within or flanking genes in the flowering time pathway.

**Figure S10:**
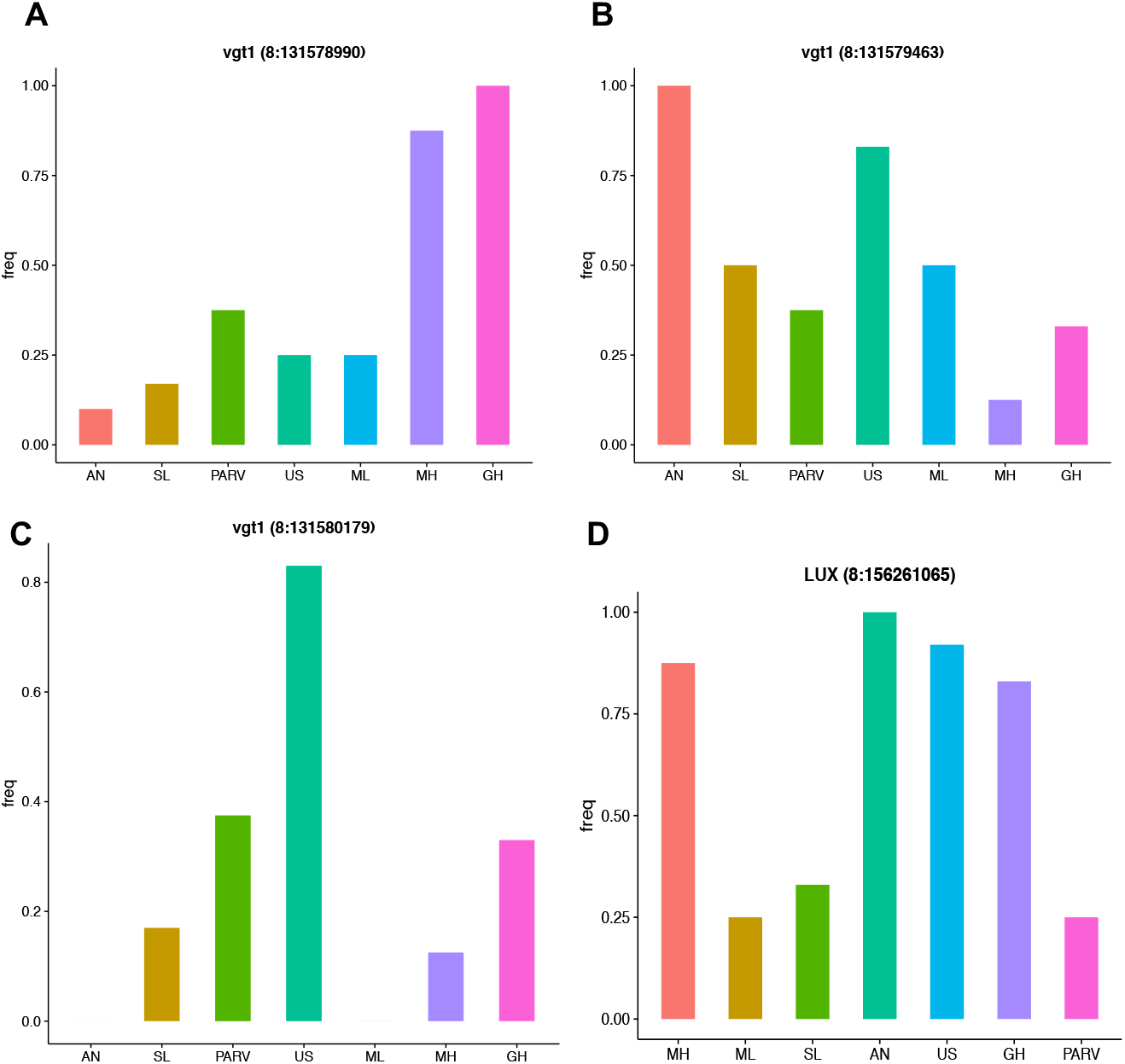
SNP frequency change in vgt1 (A, B, C) and LUX (D) among populations.

**Figure S11:**
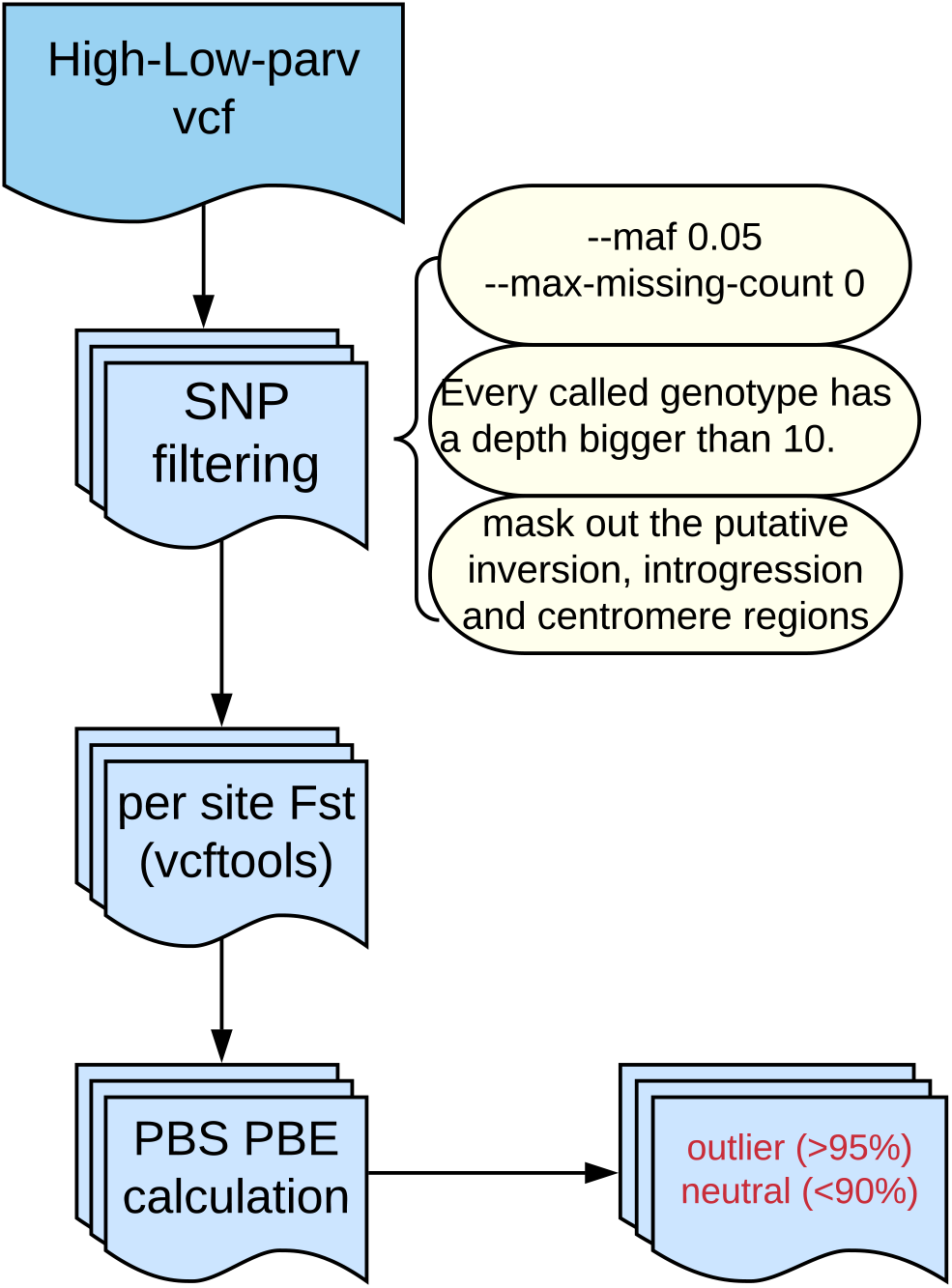
The analyses pipeline for *PBE* calculation.

**Figure S12:**
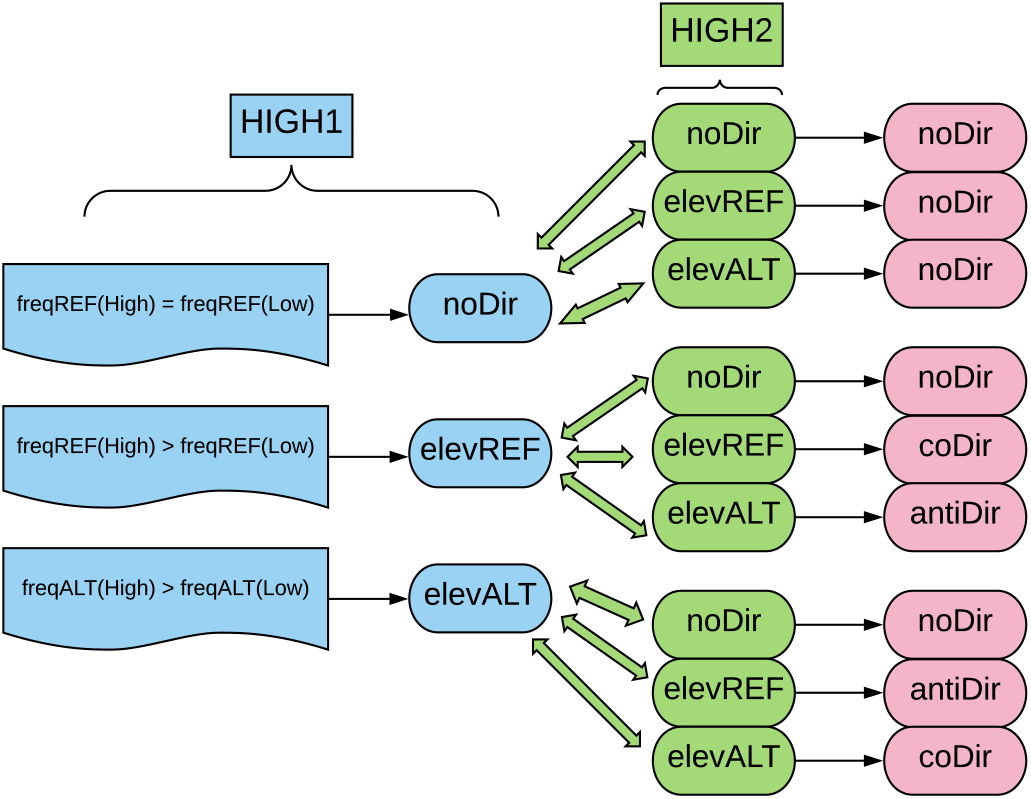
The analyses pipeline for determining co-directional and anti-directional SNPs.

**Figure S13:**
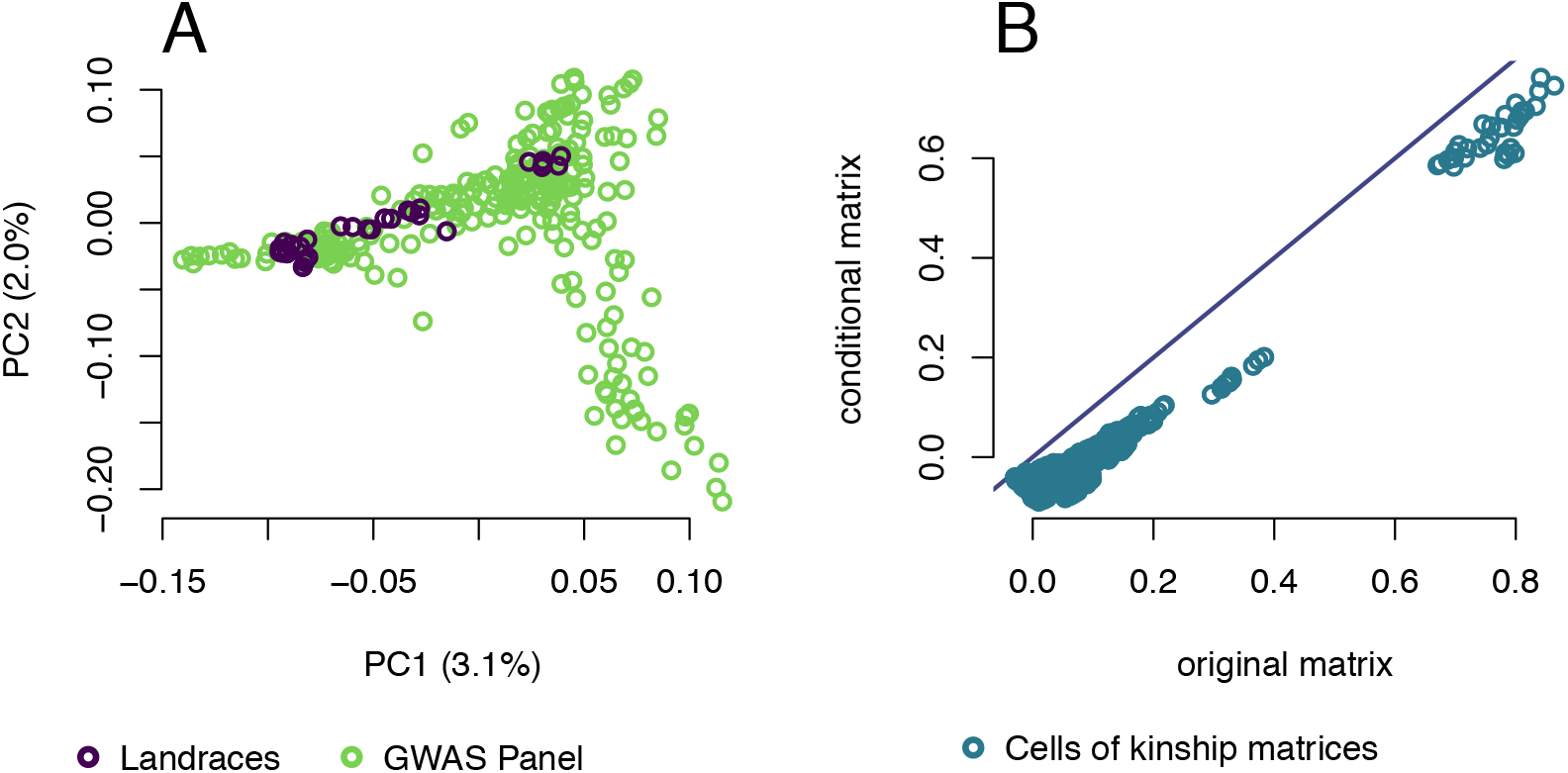
Evidence for shared population structure between the GWAS panel and the landraces. A. Individuals in both panels plotted along the first two PCs of the joint kinship matrix for the GWAS panel and the landraces. B. A comparison of all cells in the original kinship matrix for the landraces (X axis) and the conditional matrix accounting for relatedness with the GWAS panel (Y axis). The y=x line is plotted for reference.

